# Computational neural dynamics of goal-directed visual attention in macaques

**DOI:** 10.64898/2026.01.18.700191

**Authors:** Jie Zhang, Nuttida Rungratsameetaweemana, Shuo Wang

## Abstract

Goal-directed visual attention requires the dynamic integration of task goals with perceptual and mnemonic processes across distributed cortical networks. Using large-scale recordings from V4, IT, OFC, and LPFC, we identified distinct neural populations selective for attention and category. Population dynamics robustly represented visual categories during cue presentation, sustained cue information across delays, and differentiated categories and attentional states during search. Cue-related activity predicted subsequent search efficiency, linking pre-search processing to behavioral performance. The orthogonal subspace provided a crucial latent representational structure for encoding and maintaining task-relevant information across search stages. Foveal attention enhanced peripheral representations by both increasing pattern separation and reshaping representational geometry in a non-linear, context-dependent manner. Search dynamics further reflected fixation history and target detection, which modulated both response strength and representational structure. Finally, V4 and IT encoded the spatial geometry of the search array, preserving its layout. Together, these findings highlight population-level dynamics as critical mechanisms supporting goal-directed visual search.

## Introduction

Visual attention and object recognition are fundamental cognitive processes that enable primates to efficiently process and interpret their visual environment. Visual attention serves as a gateway, enhancing the perception of relevant stimuli while filtering out distractions, allowing the brain to selectively prioritize information for further processing ^1–3^. Extensive research has delineated the neural circuits underlying attentional selection, including the prefrontal, parietal, and temporal cortices, which dynamically coordinate sensory processing based on task demands ^4–9^. Among these, neurons in the inferotemporal (IT) cortex and area V4 play a critical role in integrating attention with object recognition, exhibiting attention modulation in their response properties ^10–14^, synchrony with other brain regions ^7,14–17^, and enhanced stimulus representation ^17,18^. On the other hand, visual object recognition, the ability to identify and categorize visual objects, relies on the integration of multiple visual features such as shape, color, and texture. It relies on intricate neural circuits, which allow the brain to identify and classify objects based on various visual attributes ^19^. In particular, IT neurons play a central role in the representation and processing of visual objects, demonstrating strong category selectivity ^19–21^. Neurons dedicated to face processing are found in the IT cortex and orbitofrontal cortex (OFC) ^22^, and a map of object space exists in the macaque IT cortex ^23^. Furthermore, IT neurons exhibit robust visually selective responses even in complex natural scenes, highlighting their role in recognizing objects under varying conditions ^10^. These properties position IT and V4 as critical sites for investigating how attention dynamically interacts with category-selective representations.

Visual search engages multiple interacting processes, including working memory, attentional guidance, and decision making ^24^. Yet, most prior research has examined these processes in isolation, focusing on single components rather than their dynamic interplay. Consequently, the predictive relationships between successive search events—how neural or behavioral states at one stage influence subsequent processing—remain poorly understood. Furthermore, neurophysiological studies have largely centered on single-unit responses ^1–3^; however, the population-level computations that support dynamic information processing during visual search are much less clear. In particular, how coordinated neural population activity gives rise to flexible, goal-directed behavior remains an open question. In addition, traditional analyses emphasizing firing rate changes (i.e., representations along firing rate axes) overlook the geometric structure of population activity. Beyond rate-based dimensions, information may also be encoded in the *orthogonal subspace* of the neural state space ^25^—a representational domain that has rarely been explored in visual search. Characterizing this high-dimensional geometry may offer new insight into how distributed neural populations dynamically encode and transform information to guide perception, attention, and decision making.

Recent advances in computational and systems neuroscience have illuminated the neural population dynamics underlying such computations. The dynamical systems perspective is prevalent with the motor regions serving as the most natural testing ground since time-varying behavior can be a direct output ^26–28^. However, dynamic activity is important in non-motor areas as well ^29–35^. For instance, Bayesian inference—a core principle of cognition—has been linked to latent cortical dynamics in which prior knowledge warps low-dimensional manifolds, sculpting neural trajectories to optimize behavior under uncertainty ^36^. In particular, although most evidence comes from tasks outside visual search, the neural processes supporting it are increasingly understood not as isolated tuning of single neurons but as trajectories evolving within high-dimensional state spaces. It has been shown that in the prefrontal cortex, sequence working memory relies on a geometric organization of neural dynamics, whereby complex representations can be decomposed into low-dimensional subspaces encoding ordinal and spatial information ^37^. Dynamic analyses have revealed how neural encoding of categories supports cognitive tasks with varying demands, showing that in tasks with explicit working memory requirements, population activity within each category converges toward a stable category state, resulting in binary-like categorical encoding ^38^. Furthermore, a negative correlation between the variability of V4 neurons and both attention and learning lies within a relatively small subspace of the neural state space ^39^. Notably, an orthogonal subspace of the neural state space encoded critical information, forming curved population-response manifolds that flexibly rotated depending on task context, thereby providing a representational geometry that generalizes across decision circuits ^25^. In addition, recurrent dynamics in the prefrontal state space support context-dependent computations ^40^, and a new dimensionality reduction method, latent circuit inference, has further revealed a suppression mechanism whereby contextual representations inhibit irrelevant sensory responses ^41^.

In this study, we trained macaques to perform a free-gaze visual search task using natural face and object stimuli and systematically investigated the neural dynamics underlying visual search across multiple processes. We recorded neuronal activity from a large number of units across multiple cortical regions involved in attention and object processing. Despite the well-established roles of attention and object representation that converge in areas V4 and IT—where feature-based attention modulates visual search behavior ^42^ and predicts target detection efficiency ^43^—the precise interactions and underlying neural computations between these processes remain unclear. Here, we first identified distinct populations of attention-selective and category-selective units and characterized how they dynamically contribute to goal-directed search. We analyzed population dynamics across recorded units to characterize how visual categories and attentional states are represented during cue presentation, maintained across delays, and integrated during search. We then examined how cue-related activity predicts subsequent search behavior, how foveal attention modulates peripheral representations and the geometry of neural population activity, and how fixation history and target detection influence representational structure. In particular, we examined how task-relevant information is encoded and maintained within the orthogonal subspace—a latent representational structure independent of overall firing rate changes. Finally, we assessed how spatial information about the search array is encoded in V4 and IT. Together, these analyses provide a comprehensive computational framework to understand the population-level neural mechanisms that support flexible, goal-directed visual search.

## Results

### Behavior

Two monkeys performed a free-gaze visual search task with mapped receptive fields (see **Methods** and unit summary below). Their goal was to fixate on one of two search targets that matched the category of a preceding cue (**Fig. 1A**). Each trial began with a central fixation point presented for 400 ms, followed by a cue lasting 500–1300 ms. After a 500 ms delay, a search array of 11 items appeared, which included two category-matching targets randomly positioned among 20 possible locations ^14,16,17,44^. The monkeys had up to 4000 ms to locate one of the targets and were required to maintain fixation on it for 800 ms to receive a juice reward. Fixating on either target completed the trial; the second target was not searched for. A new trial began immediately after the reward. Importantly, although both targets matched the cue category, they were always different images. During the cue and delay periods, the monkeys were required to maintain fixation.

**Fig. 1.**
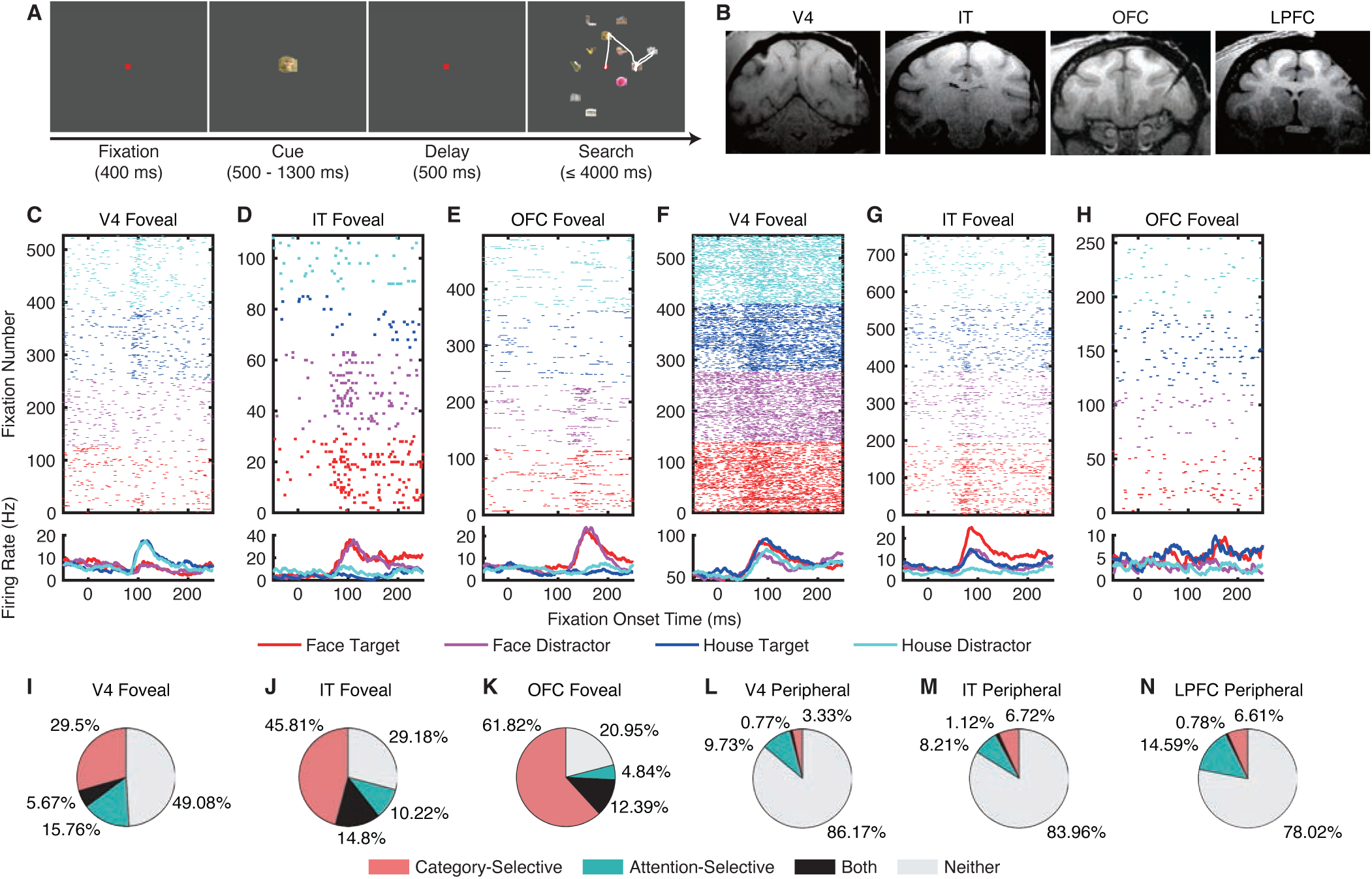
Category-selective and attention-selective units. (**A**) Task. Monkeys initiated the trial by fixating on a central point for 400 ms. A cue was then presented for 500 to 1300 ms. After a delay of 500 ms, the search array with 11 items appeared. Monkeys were required to fixate on one of the two search targets that belonged to the same category as the cue for at least 800 ms to receive a juice reward. The white trace indicates eye gazes. **(B)** MRI images show the typical recording regions of V4, TEO, TE, and the orbitofrontal cortex (OFC). **(C–E)** Example category-selective units. **(F–H)** Example attention-selective units. **(C, F)** V4 foveal units. **(D, G)** IT foveal units. **(E, H)** OFC foveal units. Each row represents a fixation. Time 0 denotes the onset of each fixation. Average firing rates are shown below. **(I–N)** Population summary of category-selective and attention-selective units.

Detailed behavioral analyses have been reported in our previous studies ^14,16,17,44^. Briefly, both monkeys performed the task at a high level of proficiency, achieving accuracy rates of 91.78% ± 0.19% (mean ± SD across sessions) for monkey S and 85.23% ± 0.41% for monkey E. The average reaction time (RT), measured from the onset of the search array to the onset of the final fixation, was 411.47 ± 67.01 ms (mean ± SD across sessions), and the mean fixation duration was 208.24 ± 153.77 ms (mean ± SD across fixations). In 13.46% ± 7.00% of correct trials, the target was located following a return fixation. Critically, the monkeys were trained to search for faces and houses that matched the category of the cue but were represented by different images. This design enabled us to probe visual attention with a diverse stimulus set while also examining mechanisms of visual object processing.

### Two distinct populations of units: category-selective and attention-selective units

We recorded a total of 6871 units from area V4, 6694 units from TEO, 1947 units from TE, 5622 units from the OFC, and 9916 units from the LPFC (**Fig. 1B**; see ^14,16,17,44^ for detailed characterization of the recording sites). Of these, 5070 units in V4, 3800 in TEO, 1251 in TE, 1470 in the OFC, and 2997 in the LPFC exhibited a significant visually evoked response (i.e., responses to the cue or search array were significantly greater than baseline; Wilcoxon rank-sum test, P < 0.05). For analyses, we combined TE and TEO into a single inferotemporal (IT) region. Foveal and peripheral receptive fields (RFs) were mapped using the visual search task and an additional visually guided saccade task (**Methods**; see also ^17^ for response consistency across tasks). Among these visually responsive units, 1624 units from V4, 1419 units from IT, 888 units from the OFC, and 32 units from the LPFC had a focal foveal RF, while 781 units from V4, 268 units from IT, no units from the OFC, and 514 units from the LPFC had a localized peripheral RF (the remaining units had broad foveal RFs or unlocalized peripheral RFs; see **Methods**).

We found that a substantial population of foveal units in V4 (35.16%, binomial P < 10^−20^; see **Fig. 1C** and **Fig. S1A, B** for examples; see **Fig. 1I** for group results), IT (60.61%, binomial P < 10^−20^; **Fig. 1D**; **Fig. S1C, D**; **Fig. 1J**), and OFC (74.21%, binomial P < 10^−20^; **Fig. 1E**; **Fig. S1E, F**; **Fig. 1K**) exhibited category selectivity, i.e., they differentiated fixations on faces versus houses (see **Methods**). We also observed category selectivity in LPFC peripheral units (7.39%, binomial P = 0.0072; **Fig. S1K, L**; **Fig. 1N**). By comparison, category selectivity in peripheral units was weak or not above chance in V4 (4.1%, binomial P = 0.86; **Fig. S1G, H**; **Fig. 1L**) and IT (7.84%, binomial P = 0.016; **Fig. S1I, J**; **Fig. 1M**). In contrast, a substantial proportion of foveal units in V4 (21.43%, binomial P = 6.65×10^−37^; see **Fig. 1F** and **Fig. S1M, N** for examples; see **Fig. 1I** for group results), IT (25.02%, binomial P < 10^−20^; **Fig. 1G**;

**Fig. S1O, P**; **Fig. 1J**) and OFC (17.23%, binomial P < 10^−20^; **Fig. 1H**; **Fig. S1Q, R**; **Fig. 1K**) were attention-selective, i.e., they differentiated fixations on targets versus distractors (see **Methods**). Peripheral units in V4 (10.5%, binomial P = 1.75×10^−10^; **Fig. S1S, T**; **Fig. 1L**), IT (9.33%, binomial P = 0.0011; **Fig. S1U, V**; **Fig. 1M**), and LPFC (15.37%, binomial P < 10^−20^; **Fig. S1W, X**; **Fig. 1N**) also demonstrated attention selectivity.

Notably, there was an interaction between category selectivity and attention selectivity in V4 foveal units (see **Fig. S1M, N** for examples and see **Fig. 1I** for group results), but this was not observed in other groups of units (**Fig. 1J–N**). Specifically, in V4, category-selective units were more likely to also be attention-selective (i.e., the proportion of attention-selective units within category-selective units [*n*_Both_ / *n*_Category-Selective_] was significantly higher than the proportion of attention-selective units within all units [*n*_Attentive-Selective_ / *n*_All_]; χ^2^-test: P = 0.0063; Bonferroni correction for comparisons in multiple brain areas; **Fig. 1I**). In contrast, this relationship was not significant for IT foveal units (P = 0.75; **Fig. 1J**), OFC foveal units (P = 0.78; **Fig. 1K**), or any peripheral units (all Ps > 0.05; **Fig. 1L–N**). Together, these results demonstrate a significant link between visual category coding and attention coding in V4 foveal units, but not in IT, OFC, or LPFC.

Lastly, we showed that the hyperplane separating fixations on faces versus houses (i.e., category selectivity) was largely orthogonal (V4: 90.18°; IT: 93.19°; OFC: 97.71°) to the hyperplane separating fixations on targets versus distractors (i.e., attention selectivity). These values did not differ from permutations except for OFC (V4: 90.01°; IT: 90.07°; OFC: 90.09°; permutation test: V4: P = 0.50; IT: P = 0.30; OFC: P < 0.001), suggesting that these attributes were encoded largely independently. These results further indicate that largely distinct populations of units encode visual categories and attention.

### Neural population dynamics during cue maintenance and search

We next quantified the dynamics of neural population activity across different stages of the visual search process (see **Methods**). We applied demixed principal component analysis (dPCA) to construct a low-dimensional neural state space and projected the population activity into it. We performed statistical comparisons using permutation tests. We focused on the foveal units in this analysis.

During search, while the average population activity of V4 foveal units did not strongly differentiate visual categories or attentional states (**Fig. 2A**), the population dynamics significantly differentiated both (**Fig. 2B**; see **Fig. S2A–D** for statistics). In IT foveal units, the average population activity exhibited a strong distinction between visual categories and also differentiated attentional states (**Fig. 2C**), which was reflected as distinct neural dynamics in state space (**Fig. 2D**; **Fig. S2A–D**). The average population activity of OFC foveal units primarily showed elevated responses to faces and attentional enhancement for fixations on faces (**Fig. 2E**). Accordingly, in neural state space, we observed a large and significant separation between visual categories and between fixations on face targets and face distractors (**Fig. 2F**; **Fig. S2A–D**).

**Fig. 2.**
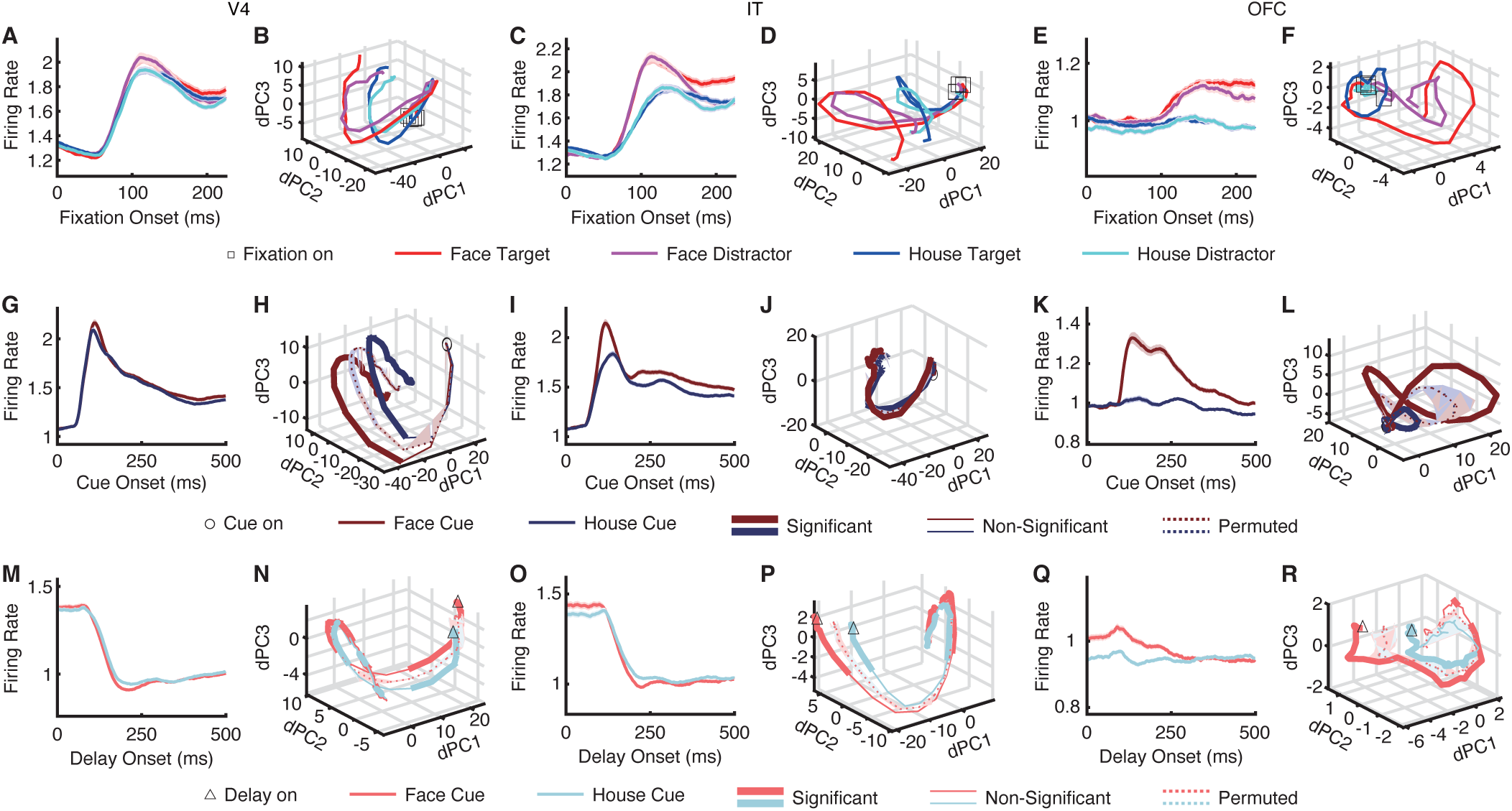
Neural population dynamics during cue maintenance and search. (**A–F**) Search. **(G–L)** Cue. **(M–R)** Delay. **(A, B, G, H, M, N)** V4 foveal units. **(C, D, I, J, O, P)** IT foveal units. **(E, F, K, L, Q, R)** OFC foveal units. **(A, C, E, G, I, K, M, O, Q)** Mean normalized firing rate. Error shades denote ±SEM across units. **(B, D, F, H, J, L, N, P, R)** Neural population dynamics. Thick lines indicate time points with significant differences between conditions (permutation test of the separation of the curves: P < 0.05, corrected by false discovery rate [FDR] across time points ^61^), while thin lines indicate non-significant time points. Solid lines represent observed data, and dotted lines represent permuted data. Shaded areas around dotted lines indicate ±SD across permutation runs (*n* = 1000).

During cue presentation (note that there was no attention component), while the average population activity of V4 foveal units barely showed differences between faces and houses (**Fig. 2G**), the neural population dynamics in state space clearly and significantly differentiated them (**Fig. 2H**), suggesting that this population encoded visual category information. IT foveal units (**Fig. 2I**) and OFC foveal units (**Fig. 2K**) differentiated faces from houses, and their neural dynamics correspondingly diverged in state space (**Fig. 2J, L**). Interestingly, during cue maintenance, the average population activity of V4 (**Fig. 2M**) and IT (**Fig. 2O**) foveal units barely differentiated faces and houses. However, the neural dynamics significantly differentiated them (**Fig. 2N, P**), suggesting that cue information was maintained through the delay period. Similar results were observed for OFC foveal units (**Fig. 2Q, R**). Notably, we replicated these findings in a separate identity-matching task, in which monkeys searched for an identical target to the cue (unlike the category-matching task above, the identity-matching task included only one search target; **Fig. S2E, F**).

Together, these results demonstrate that neural population dynamics, rather than average firing rates, robustly encode and maintain task-relevant information across stages of visual search. While V4, IT, and OFC foveal units exhibited limited category– or attention-related modulation at the level of mean activity, their population dynamics revealed clear representations of visual categories during cue presentation, sustained maintenance of cue information during the delay, and strong differentiation of both visual categories and attentional states during search. These findings highlight the critical role of population-level temporal structure in supporting flexible, goal-directed behavior.

### Neural activity during cue presentation and maintenance predicts search efficiency

We next investigated whether neural activity before search onset influenced and predicted subsequent search efficiency. We sorted all trials by the total number of fixations required to complete each trial.

In V4, we found a substantial population of units whose activity during cue presentation and maintenance was significantly correlated with the number of fixations required to complete the trial (36.04%, binomial P < 10^−20^; see **Fig. 3A, B** for examples and **Fig. 3C–E** for group summary; note that the units were selected based on mean activity from 0 to 200 ms after *delay* onset, but qualitatively the same results were obtained when using mean activity from 0 to 500 ms after *cue* onset). These units comprised two subgroups: one showing a positive correlation with the number of fixations (i.e., lower search efficiency; 24.65%; **Fig. 3A, C, D**; Pearson correlation: cue: *r*(5) = 0.91, P = 0.03; delay: *r*(5) = 0.92, P = 0.03) and another showing a negative correlation (i.e., higher search efficiency; 11.39%; **Fig. 3B, C, E**; cue: *r*(5) = −0.96, P = 0.01; delay: *r*(5) = −0.97, P = 0.008). These correlations held for both face and house cues (**Fig. 3D**). Because the majority of units were positively correlated with the number of fixations required to complete the search, this suggests that lower neural activity during cue processing and maintenance was associated with greater search efficiency.

**Fig. 3.**
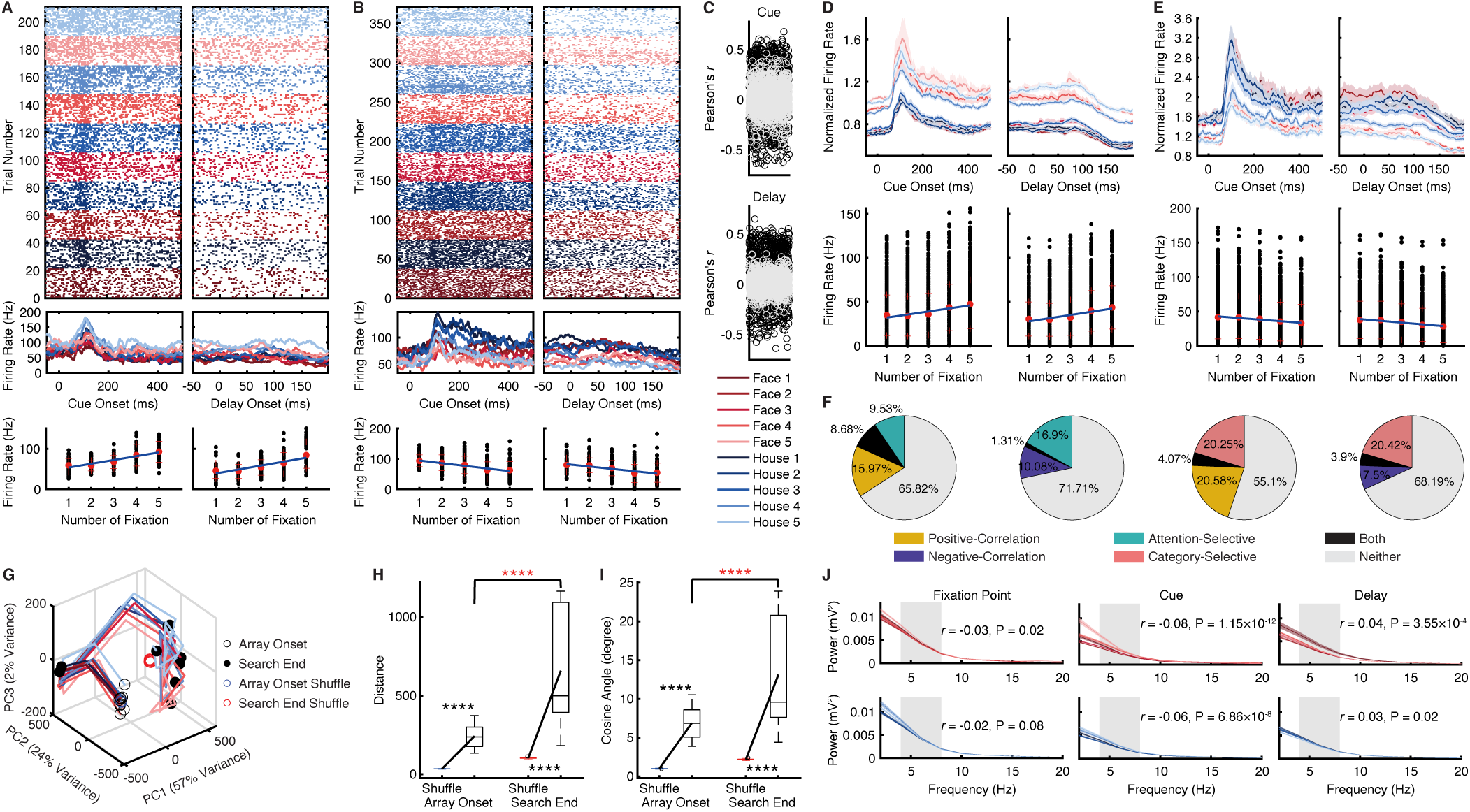
Neural activity in V4 during cue presentation and maintenance predicts search efficiency. (**A**) An example unit exhibiting higher activity for trials requiring more fixations to complete (i.e., positive correlation). **(B)** An example unit exhibiting higher activity for trials requiring fewer fixations to complete (i.e., negative correlation). (Top) Neural activity during cue presentation and maintenance, sorted by the number of fixations required to complete the search. (Middle) Mean firing rate. (Bottom) Correlation between average neural activity and the number of fixations needed to complete the search. Each dot represents a trial. The central red dot represents the mean, and the red crosses denote ±SD. The line represents the linear fit. Red: face cues. Blue: house cues. **(C)** Summary of the correlation coefficient (Pearson’s *r*) between average neural activity and the number of fixations required to complete the search. Each circle represents a unit. Black: units with a significant correlation. Gray: units with a non-significant correlation. **(D)** Group summary of units showing a positive correlation between activity and the number of fixations required during the search. **(E)** Group summary of units showing a negative correlation between activity and the number of fixations required during the search. (Upper) Group PSTH. Error shading denotes ±SEM across units. (Lower) Correlation (cue: mean activity from 0 to 500 ms after cue onset; delay: mean activity from 0 to 200 ms after delay onset). Each dot represents a unit. The central red dot represents the mean, and the red crosses denote ±SD. The line represents the linear fit. **(F)** Overlap between units showing significant correlations and attention-selective or category-selective units. Shown are units selected during the delay period (mean activity from 0 to 200 ms after delay onset). **(G)** Neural population dynamics. The open black circle denotes the onset of the search (array onset), and the solid black circle denotes its completion. The blue and red circles indicate permuted data at the search onset and end, respectively. **(H)** Representational distance for the population of units. **(I)** Angle between neuronal vectors. In each box, the central mark indicates the median, the edges represent the 25th and 75th percentiles, the whiskers extend to the most extreme data points considered not to be outliers by the algorithm, and outliers are plotted individually. Black asterisks indicate a signifiant difference between the observed and permuted data. Red asterisks indicate a significant difference between search onset and end using a two-tailed paired *t*-test. ****: P < 0.0001. **(J)** Power as a function of the number of fixations required to complete the search. The gray shaded area indicates the theta frequency band (4–12 Hz) used to calculate the correlation between power and the number of fixations.

We found that the neuronal population showing a positive correlation with the number of fixations significantly overlapped with both the attention-selective units ([*n*_Both_ / *n*_Positive-Correlation_] versus [*n*_Attentive-Selective_ / *n*_All_]; χ^2^-test: P < 10^−10^; **Fig. 3F**) and the category-selective units ([*n*_Both_ / *n*_Positive-Correlation_] versus [*n*_Category-Selective_ / *n*_All_]; P = 5.62×10^−5^). Similarly, the neuronal population showing a negative correlation with the number of fixations also significantly overlapped with both the attention-selective units ([*n*_Both_ / *n*_Negative-Correlation_] versus [*n*_Attentive-Selective_ / *n*_All_]; P = 0.006) and the category-selective units ([*n*_Both_ / *n*_Negative-Correlation_] versus [*n*_Category-Selective_ / *n*_All_]; P = 4.10×10^−4^). This result suggests that search efficiency is linked, at least in part, to the representation of attentional and categorical information, indicating a partially integrated coding scheme.

Importantly, we observed that differential cue processing led to distinct neural dynamics during search (**Fig. 3G**), and that the neural representation of the search target (i.e., the search cue) diverged even further by the end of the search (**Fig. 3H, I**; two-tailed paired *t*-test: representational distance: *t*(44) = 7.80, P = 7.88×10^−10^, *d* = 0.39, 95% CI = [31.72, 246.66]; angle between neuronal vectors: *t*(44) = 5.56, P = 1.49×10^−6^, *d* = 0.63, 95% CI = [2.45, 6.97]), suggesting that early cue-related processing shapes the evolving population code during visual search.

To explore the underlying mechanism, we conducted a frequency analysis. Interestingly, we found that more efficient search (i.e., fewer fixations) was associated with enhanced theta-band oscillations during cue presentation (**Fig. 3J**; Pearson correlation: both Ps < 10^−7^), but with reduced theta-band oscillations during cue maintenance (**Fig. 3J**; both Ps < 0.05), suggesting that theta activity during cue processing may modulate attentional engagement and improve subsequent search performance.

Notably, we found similar results for IT (**Fig. S3C, D**), OFC (**Fig. S3E**), and LPFC (**Fig. S3F**) units. Specifically, in each of these regions we identified subpopulations of units whose activity during cue presentation and maintenance was significantly correlated with the number of fixations required to complete the trial. As in V4, these correlations included both positively correlated units (reflecting lower search efficiency) and negatively correlated units (reflecting higher search efficiency), and were observed for both face and house cues. Moreover, overlapping analyses revealed that these efficiency-predicting units also intersected significantly with attention-selective and category-selective populations. Additionally, more efficient search was associated with enhanced theta-band oscillations during cue presentation but with reduced theta-band oscillations during cue maintenance. Overall, these results indicate that multiple cortical areas—including V4, IT, OFC, and LPFC—are engaged in representing neural signals predictive of search efficiency.

Lastly, we obtained similar results when analyzing foveal (**Fig. S3A, C, E**) and peripheral (**Fig. S3B, D, F**) units separately.

Together, these results suggest that across multiple cortical areas, pre-search cue processing predicts subsequent visual search efficiency and modulates neural dynamics, revealing an integrated brain network and a coordinated mechanism for attentional engagement and target selection.

### Neural processing across stages of visual search in the orthogonal subspace

While the preceding analyses based on firing rates revealed neural dynamics at each stage of search and their relationship to behavior, they did not capture the full structure of neural population activity. Recent work has shown that critical task-related information can be represented in the *orthogonal subspace* of neural state space—dimensions that are independent of overall firing rate changes ^25^. Examining how information evolves within this orthogonal subspace allows us to uncover not only latent population codes but also the transitions between stages of visual search, including cue encoding, delay maintenance, and target detection.

Indeed, in V4 units non-selective for attention (**Fig. 4A, C**), activity along the average firing rate (parallel) axis did not distinguish fixations on targets versus distractors over time, as expected. In contrast, activity in the orthogonal plane reliably differentiated targets from distractors across time, emerging even before fixation onset and persisting throughout the fixation (**Fig. 4A, C**). A similar pattern was observed in IT units non-selective for attention (**Fig. 4C**), particularly during later time windows in the orthogonal subspace. Likewise, in V4 units non-selective for category (**Fig. 4B, D**), the parallel axis failed to separate fixations on faces versus houses, whereas activity in the orthogonal plane robustly distinguished the two categories across time (**Fig. 4B, D**). This effect was also evident in IT units non-selective for category (**Fig. 4D**). Together, these results indicate that the orthogonal subspace captures additional neural processing not reflected in mean firing rates.

**Fig. 4.**
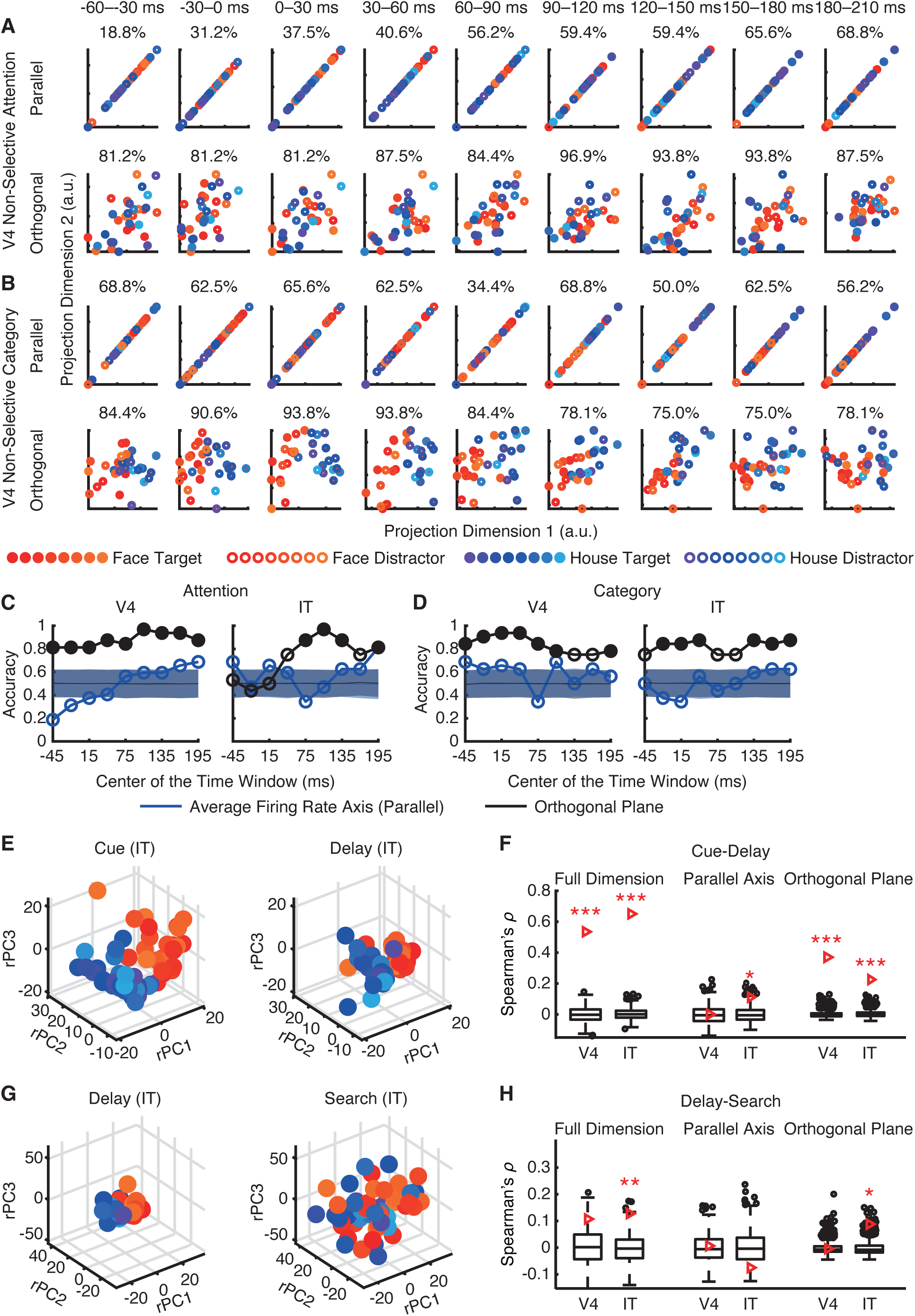
Neural processing across stages of visual search in the orthogonal subspace. **(A)** Projection of the population response of V4 non-selective units onto the attention subspace across time. **(B)** Projection of the population response of V4 non-selective units onto the category subspace across time. (Upper) Projection onto the average firing rate axis; (Lower) Projection onto the plan orthogonal to the average firing rate axis. Classification accuracy is indicated in the title. **(C)** Classification accuracy for attention. **(D)** Classification accuracy for category. **(E)** Projection of the population response of IT foveal units during cue and delay. **(F)** Correlation between RDMs during cue and delay. **(G)** Projection of the population response of IT foveal units during delay and search. **(H)** Correlation between RDMs during delay and search. The central mark represents the median of the null distribution, the top and bottom edges correspond to the 75th and 25th percentiles, respectively, and circles indicate outliers. The red triangle indicates the observed value. Asterisks indicate significant correlations (permutation test, *: P < 0.05, **: P < 0.01, and ***: P < 0.001).

In addition, consistent patterns were observed in face-selective (**Fig. S4A, C, E, F**) and house-selective (**Fig. S4B, D, G, H**) units. Specifically, both face-selective and house-selective units in V4 and IT differentiated faces from houses along the average firing rate axis (**Fig. S4F, H**; particularly in later time windows), as expected, but did not encode attention along this axis (**Fig. S4E, G**), consistent with largely distinct neuronal populations for category and attention coding. In contrast, activity in the orthogonal plane not only distinguished visual categories (**Fig. S4F, H**) but also captured attentional modulation (**Fig. S4E, G**), especially in face-selective units. Thus, the orthogonal subspace again revealed additional information processing not evident in mean firing rates.

Importantly, by analyzing stimulus representations across different stages of the search (**Fig. 4E–H**; see **Methods**), we found that cue information was maintained throughout the delay period (**Fig. 4E, F**), primarily within the orthogonal plane rather than along the parallel axis (**Fig. 4F**). This pattern was consistent across both V4 and IT units (**Fig. 4F**; permutation test: both Ps < 0.001). Moreover, stimulus information encoded during the delay was further transmitted to the search period in IT units (**Fig. 4G, H**), again predominantly through the orthogonal subspace (P = 0.014).

Together, these findings demonstrate that the orthogonal subspace provides a crucial latent representational structure through which the brain encodes and maintains task-relevant information across stages of visual search, supporting continuous information processing beyond what can be captured by mean firing rates alone.

### Attention modulation of neural representations in the peripheral visual fields

Peripheral units are important for saccade selection ^16,42^. What is the interaction between foveal and peripheral processing, and how does attention modulate peripheral processing? To address these questions, we next investigated the neural representations of peripheral units as a function of foveal content and attentional state (see **Methods**). We focused on non-category-selective peripheral units in each brain area to isolate effects driven by attention modulation rather than intrinsic category tuning or baseline category preferences.

First, we found that foveal attention enhanced neural pattern separation in V4 peripheral units (**Fig. 5A–D**). The neural representational distance between all stimuli (including four categories of distractors and two categories of targets in the peripheral RF) was significantly greater when monkeys fixated on targets versus distractors (**Fig. 5C**; two-tailed paired *t*-test: *t*(14) = 6.17, P = 2.44×10^−5^, *d* = 1.59, 95% CI = [72.08, 148.95]), and the angle between neuronal vectors was also significantly larger (**Fig. 5D**; *t*(14) = 10.53, P = 4.94×10^−8^, *d* = 2.72, 95% CI = [22.86, 34.57]). Interestingly, when monkeys fixated on targets (**Fig. 5A**), neural representations of distractors were similar across all four distractor categories, whereas representations of targets were substantially enhanced, primarily driving the increased pattern separation. In contrast, when monkeys fixated on distractors (**Fig. 5B**), neural representations of targets were less distinct, while face and house distractors were more distinct than the other two distractor types, indicating more graded pattern separation across stimulus categories. Thus, beyond the general enhancement of peripheral pattern separation by foveal attention, these results reveal nuanced, non-linear mechanisms of attentional modulation.

**Fig. 5.**
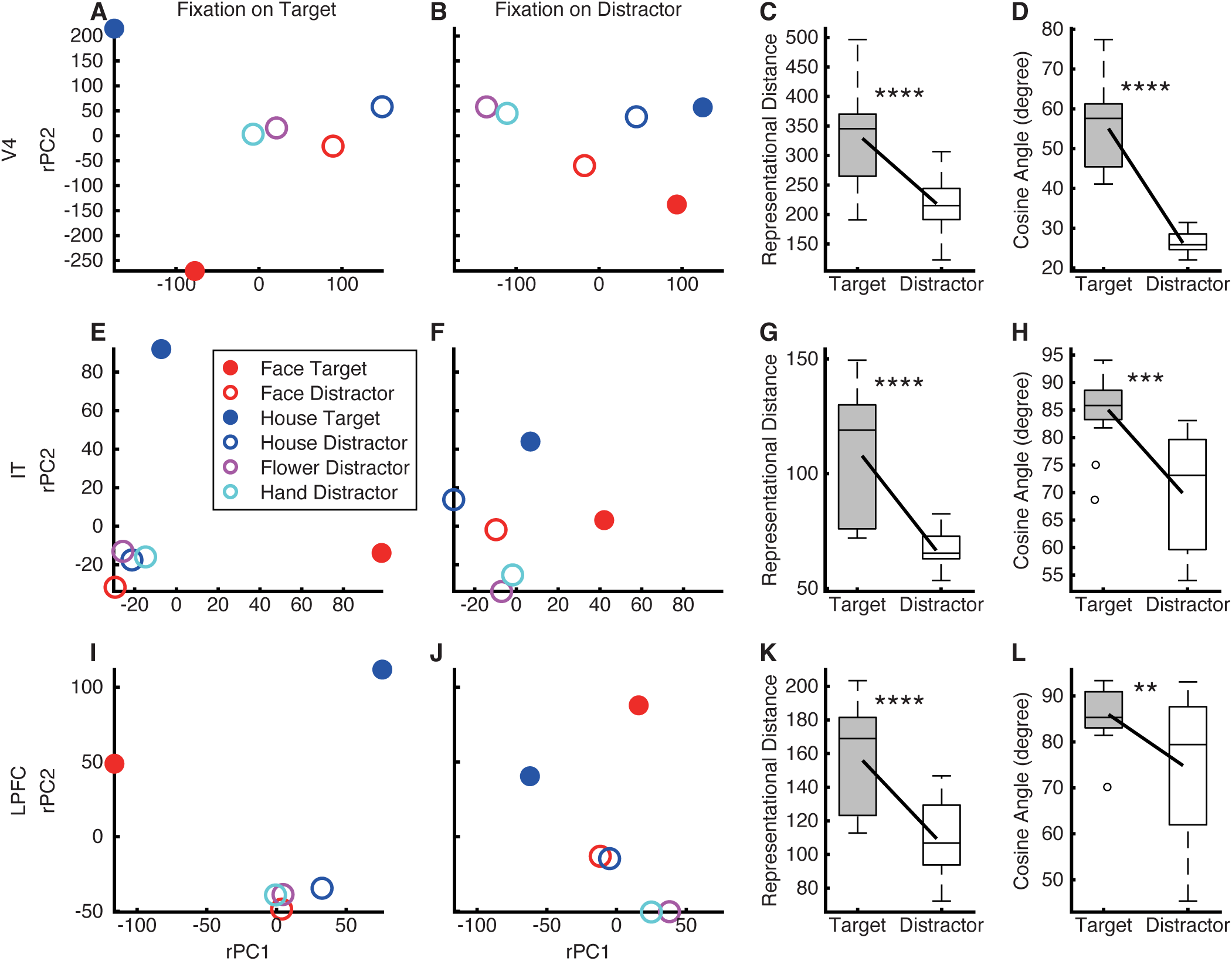
Attention modulation of neural representations in the peripheral visual fields. (**A–D**) V4 units. **(E–H)** IT units. **(I–L)** LPFC units. All units are non-category-selective peripheral units. **(A, B, E, F, I, J)** The population response for each stimulus with specific attention-category combination projected to the corresponding foveal attentional state subspace. Responses were obtained through linear regression of averaged activity. Dimensions of the neural state subspaces are represented by the rotated principal components (rPCs). **(A, E, I)** Fixations on targets. **(B, F, J)** Fixations on distractors. **(C, G, K)** Representational distance for the population of units. **(D, H, L)** Angle between neuronal vectors. In each box, the central mark indicates the median, the edges represent the 25th and 75th percentiles, the whiskers extend to the most extreme data points considered not to be outliers by the algorithm, and outliers are plotted individually. Asterisks indicate a significant difference between subspaces (distractor versus target) using a two-tailed paired *t*-test. *: P < 0.05, **: P < 0.01, ***: P < 0.001, and ****: P < 0.0001.

Similarly, both IT peripheral units (**Fig. 5E–H**; representational distance: *t*(14) = 6.87, P = 7.7×10^−6^, *d* = 1.77, 95% CI = [28.36, 54.12]; angle: *t*(14) = 4.92, P = 2.25×10^−4^, *d* = 1.27, 95% CI = [8.52, 21.69]) and LPFC peripheral units (**Fig. 5I–L**; representational distance: *t*(14) = 7.57, P = 2.58×10^−6^, *d* = 1.96, 95% CI = [33.31, 59.64]; angle: *t*(14) = 3.08, P = 8.2×10^−3^, *d* = 0.79, 95% CI = [3.37, 18.85]) exhibited enhanced pattern separation by foveal attention. Importantly, foveal attention modulated neural representations in a similarly non-linear manner as in V4: when monkeys fixated on targets (**Fig. 5E, I**), representations of targets were strongly enhanced while distractors collapsed onto a common representation, whereas when fixating on distractors (**Fig. 5F, J**), target representations became less distinct and face and house distractors were more separated than others.

Together, these findings demonstrate that foveal attention enhances peripheral neural representations across visual and prefrontal areas, not only by globally increasing pattern separation but also by reshaping representational geometry in a non-linear, context-dependent manner.

### Neural dynamics change as a function of task contexts

How are neural representations dynamically modulated by task context? First, we compared first-fixated versus re-fixated search items in foveal units from each brain area. We found that re-fixation attenuated the neural response in V4 (**Fig. 6A, B**; two-tailed paired *t*-test across all units in a time window 150–250 ms after fixation onset: *t*(1622) = 9.43, P = 1.41×10^−20^, *d* = 0.23, 95% CI = [0.22, 0.33]) and IT (**Fig. 6C, D**; *t*(1417) = 6.32, P = 3.4×10^−10^, *d* = 0.17, 95% CI = [0.2, 0.38]), but not in OFC (**Fig. 6E, F**; *t*(886) = 1.82, P = 0.07, *d* = 0.06, 95% CI = [−0.003, 0.08]). Interestingly, the neural representational geometry was consistent between first fixations and re-fixations (**Fig. 6G–I**), and across brain areas, we did not observe significant differences in neural representational distance (**Fig. 6J**; all Ps > 0.05 except for V4) or in the angle between neuronal vectors (**Fig. 6K**; all Ps > 0.05). Similar results were obtained for fixations on distractors only (**Fig. S5**).

**Fig. 6.**
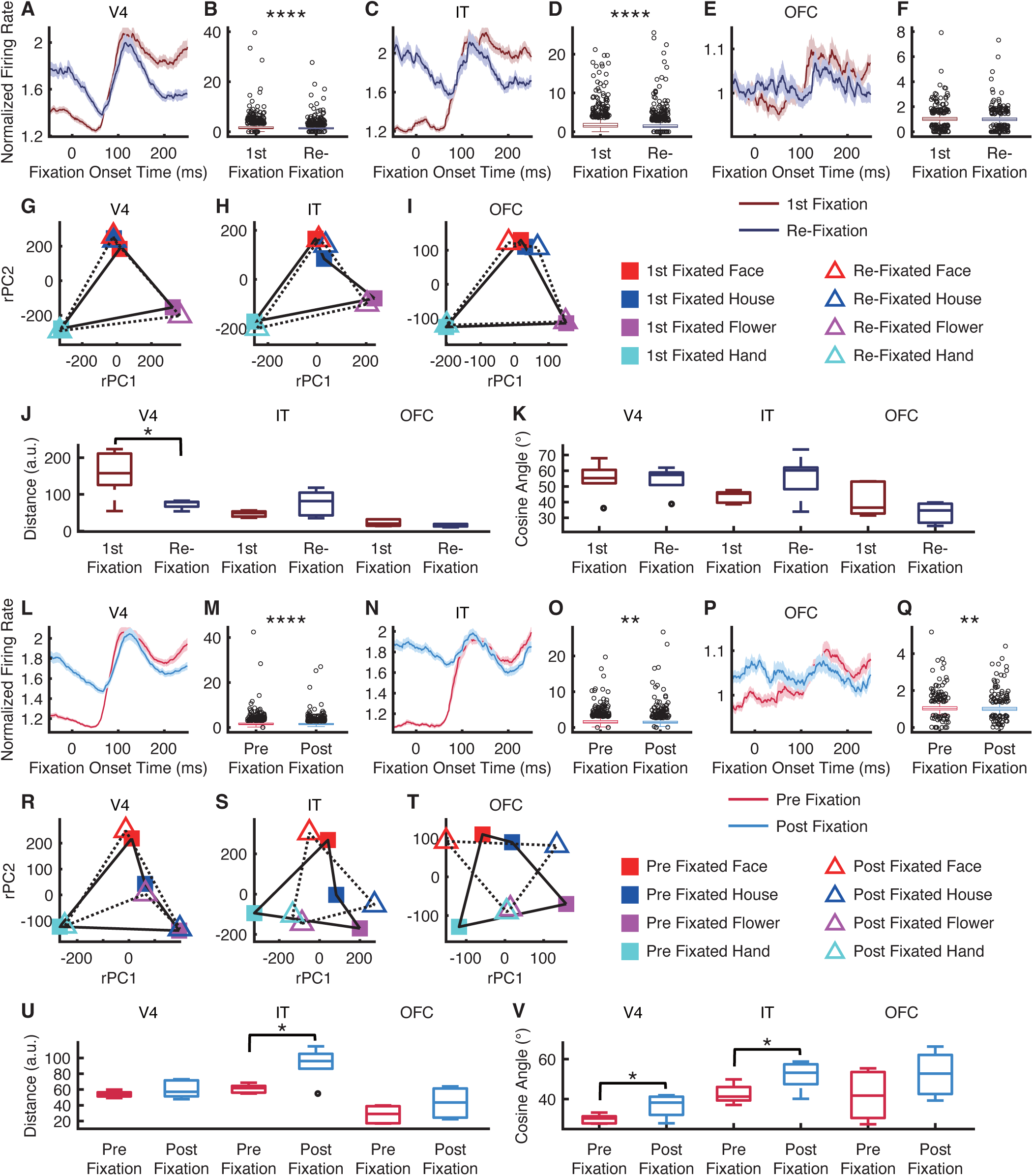
Neural dynamics change as a function of task contexts. (**A–K**) Comparison of first-fixated versus re-fixated search items. **(L–V)** Comparison of fixations on distractors before versus after the first fixation on the target. **(A, B, G, L, M, R)** V4 units. **(C, D, H, N, O, S)** IT units. **(E, F, I, P, Q, T)** OFC units. **(A, C, E, L, N, P)** Mean normalized firing rate. Error shades denote ±SEM across units. **(B, D, F, M, O, Q)** Mean normalized firing rate in a time window 150–250 ms after fixation onset. In each box, the central mark indicates the median, the edges represent the 25th and 75th percentiles, the whiskers extend to the most extreme data points considered not to be outliers by the algorithm, and outliers are plotted individually. Asterisks indicate a significant difference between conditions using a two-tailed paired *t*-test. *: P < 0.05, **: P < 0.01, ***: P < 0.001, and ****: P < 0.0001. **(G–I, R–T)** Projection of the population response onto the neural state space. **(J, U)** Representational distance for the population of units. **(K, V)** Angle between neuronal vectors.

We next investigated whether fixating on targets altered responses to distractors (e.g., ^45^). Indeed, across brain areas, responses to distractors were attenuated following the first fixation on the target (**Fig. 6L–Q**; V4: *t*(1622) = 6.64, P = 4.33×10^−11^, *d* = 0.16, 95% CI = [0.09, 0.16]; IT: *t*(1417) = 2.77, P = 0.006, *d* = 0.07, 95% CI = [0.02, 0.15]; OFC: *t*(886) = 2.58, P = 0.01, *d* = 0.09, 95% CI = [0.008, 0.059]). Notably, the neural representational geometry changed in V4 (**Fig. 6R**), IT (**Fig. 6S**) and OFC (**Fig. 6T**) across these distractor fixations. We observed an increase in neural representational distance in IT (**Fig. 6U**; *t*(5) = −3.59, P = 0.02, *d* = −1.47, 95% CI = [−53.09, −8.78]) and an increase in the angle between neuronal vectors in V4 (**Fig. 6V**; *t*(5) = −2.87, P = 0.04, *d* = −1.17, 95% CI = [−12.29, −0.67]) and IT (*t*(5) = −2.7, P = 0.04, *d* = −1.1, 95% CI = [−18.02, −0.46]).

Together, these results demonstrate that neural dynamics during search are shaped both by fixation history and by the attainment of the search target. While re-fixations attenuated responses in V4 and IT without altering the underlying representational geometry, locating the target led to a broader suppression of distractor responses and changes in neural representational geometry. These findings suggest that multiple cortical areas dynamically adjust both response strength and representational structure to optimize visual search behavior.

### Neural representation of search array spatial geometry in V4 and IT

Lastly, we investigated whether neural population activity encoded spatial information and the geometric structure of the search array (e.g., ^37^). Specifically, we examined whether the neural population exhibited a geometry in state space that mirrored the spatial distribution of the search items (**Fig. 7A**).

**Fig. 7.**
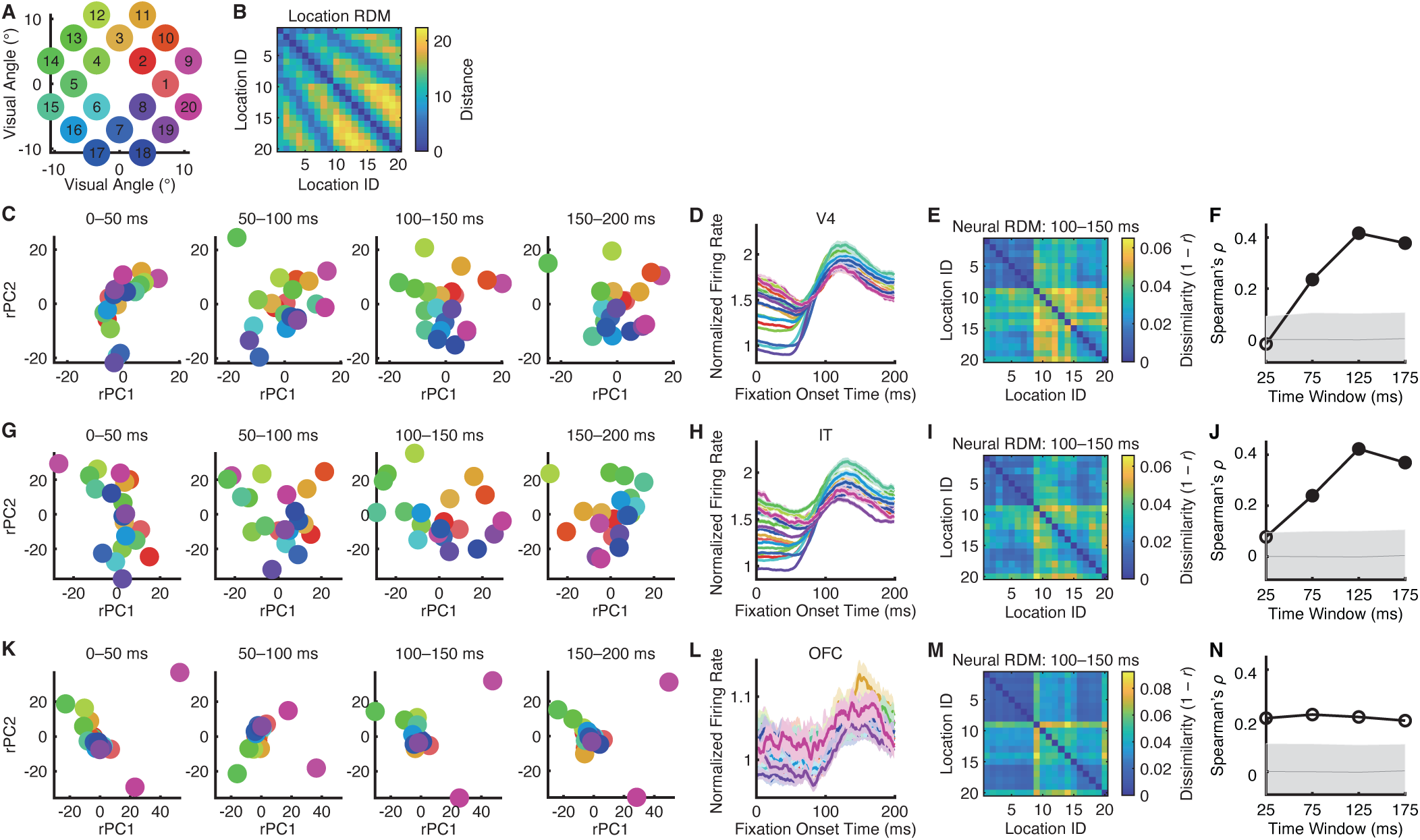
Neural representation of search array spatial geometry. **(A)** Search array locations. **(B)** Representational dissimilarity matrix (RDM) of search array locations. Color coding shows dissimilarity values (i.e., distances between physical locations). **(C–F)** V4 units. **(G–J)** IT units. **(K–N)** IT units. **(C, G, K)** Projections of neuronal vectors across four time windows (0–50 ms, 50–100 ms, 100–150 ms, and 150–200 ms) into the common neural state space. **(D, H, L)** Mean firing rate for each search array location. Shaded areas denote ±SEM across units. **(E, I, M)** Neural RDM of search array locations in the 100–150 ms time window. Color coding shows dissimilarity values (i.e., 1 − Pearson’s correlation between neuronal vectors). **(F, J, N)** Correlation between neural and physical RDMs. Solid circle: P < 0.05 (permutation test; Bonferroni-corrected across comparisons). Shaded areas denote ±SD across permutation runs.

Indeed, the mean firing rate of all V4 units differentiated search array locations and exhibited similar or clustered activity for locations adjacent to each other (**Fig. 7D**). Importantly, when we projected the neural population activity into the neural state space, we observed a spatial distribution of encoded search items that matched the actual physical search array (**Fig. 7C**), suggesting that the neural representational geometry corresponded to the physical array structure. Our RSA (see **Methods**) confirmed that the pairwise distances of neuronal vectors (**Fig. 7E**) correlated with the pairwise distances between the physical locations (**Fig. 7B**) of the search items (**Fig. 7F**). Such correspondence started in the 50–100 ms time window, peaked in the 100–150 ms window, and was sustained through the 150–200 ms window of the fixation (**Fig. 7F**). Interestingly, while the mean firing rate showed the strongest differentiation of locations in the early fixation time window (**Fig. 7D**), the later time window better encoded the full spatial layout of the search array (**Fig. 7C, F**), indicating a transition from coarse rate-based coding to a more structured population-level representation of spatial information.

Similar results were observed for IT units: the mean firing rate differentiated search array locations and exhibited clustered activity for adjacent locations (**Fig. 7H**). Projection into the neural state space revealed a spatial distribution of encoded items that resembled the actual array (**Fig. 7G**). RSA further demonstrated that, beginning in the 50–100 ms time window, the pairwise distances of neuronal vectors (**Fig. 7I**) correlated with the pairwise distances between the physical locations (**Fig. 7B**) of the search items (**Fig. 7J**), with correspondence peaking in the 100–150 ms time window (**Fig. 7G, J**). In contrast, OFC units did not encode the spatial information of the search array (**Fig. 7K–N**).

In the above analyses, we included all units; however, the effects were primarily driven by non-selective units, and we confirmed similar results when restricting the analysis to this subset. By contrast, attention– and category-selective units did not encode such geometry, suggesting that they primarily represent other task-related information.

Together, these findings demonstrate that V4 and IT populations encode the spatial layout of the search array in a manner that preserves its geometric structure, providing a potential substrate for spatial organization during visual search.

## Discussion

In this study, we systematically characterized neural coding in foveal and peripheral RFs across V4, IT, OFC, and LPFC during visual search. Across areas and RF types, we identified two largely distinct classes of units: those encoding attentional state (targets versus distractors) and those encoding the categorical identity of visual stimuli (faces versus houses). Beyond delineating the computational dynamics of each class, we demonstrated that population-level neural activity, rather than mean firing rates, robustly encoded task-relevant information across cue, delay, and search epochs. Cue-related activity predicted subsequent search efficiency, linking pre-search processing to behavioral performance. In particular, task-relevant information was encoded and maintained within the orthogonal subspace—a latent representational structure independent of overall firing rate changes. Furthermore, we uncovered a robust interaction between foveal attentional states and peripheral category representations, showing that attention directed at the fovea reshapes the representational geometry of peripheral inputs. Moreover, search dynamics further reflected fixation history and target detection, which modulated both response strength and representational structure. Lastly, V4 and IT encoded the spatial geometry of the search array, preserving its layout. Together, these findings highlight population dynamics as a fundamental mechanism for goal-directed behaviors.

We trained macaques to perform a free-gaze visual search task with naturalistic face and object stimuli, enabling a detailed investigation of how visual information is represented and how attention shapes these representations. This paradigm uniquely positions our study to examine the intricate interplay between attention coding and visual category coding across multiple cortical areas. In this study, we demonstrated both distinct and overlapping coding schemes across cortical areas and RF types, underscoring the dynamic and distributed nature of attentional modulation in visual processing. On the one hand, our results align with previous findings showing units that simultaneously encode visual features and attentional state ^17^, supporting a multidimensional representational framework ^46^. Notably, units encoding visual features exhibited distinct attentional effects and spike-LFP coherence patterns, suggesting a unique computational role in coordinating local and long-range processing during visual search ^17^. Our findings also corroborate prior work using a similar naturalistic free-gaze visual search task, which reported that V4 neurons integrate both bottom-up visual responses and top-down target-related modulation ^47^. On the other hand, our results parallel studies in the human amygdala and hippocampus, where largely independent populations of neurons encode visual categories and attentional effects in comparable visual search tasks ^45^. This divergence raises an important open question: at what stage of the visual processing hierarchy do feature-based and attentional signals become integrated or remain segregated? Our present findings provide an important first step toward understanding how attention and visual information coding interact during naturalistic search. Future work combining laminar recordings, causal perturbations, and network-level modeling will be critical for dissecting how attentional modulation is implemented across hierarchical circuits.

Computational models and theories of visual search suggest that it involves multiple interacting processes ^24^, and our previous work has revealed a network of brain regions that encode these processes ^48^, among which a working memory component is particularly critical for maintaining the target representation throughout search. Working memory functions rely on a distributed network of brain regions ^49^. For instance, neurons in the medial temporal lobe (MTL) encode the working memory of the search cue ^45^, and persistently active neurons in both the human MTL and medial frontal cortex (MFC) support working memory maintenance ^50^. Extending these findings, the present study demonstrates that neural populations across V4, IT, and OFC also encode the contents of the cue during the maintenance period (**Fig. 2**). Critically, the strength of this maintenance signal predicted search efficiency (**Fig. 3**), suggesting that sustained representations in these areas are functionally relevant for guiding visual search. These results are consistent with prior studies showing that prefrontal neurons synchronize their activity at gamma frequencies when monkeys retain multiple objects across a short delay in a working memory task ^51^. Together, our findings provide convergent evidence that visual search relies on a distributed working memory network, spanning sensory, associative, and prefrontal regions. This network not only maintains cue information but also links it to ongoing attentional and perceptual processes, thereby optimizing search efficiency.

Our data reveal that neurons encode devolving task-relevant information within the visual and prefrontal neural state spaces throughout the search process to support efficient goal-directed visual behavior. These representations exhibit a geometric structure that reflects sensory, task-relevant, and behavior-related information. Importantly, such representations emerge at the neural population level rather than from single neurons or simple linear averaging across neurons. This finding aligns with previous work showing that neural gain modulation arises only at the collective level and can be characterized by matrix-based computations in the LPFC ^37^. Furthermore, previous studies have shown that neural subspaces encode attention ^39^ and category ^38^ information. In this study, we found that these two subspaces are nearly orthogonal, with distinct neuronal populations encoding visual categories and attention. Such encoding—where different types of information are represented in orthogonal subspaces—has also been reported in studies of sequence working memory ^37^ and in neural representations that integrate sensory percepts with memories of recent stimuli. Furthermore, we observed that projections onto the orthogonal subspace captured more information about category, attention, and cue maintenance than projections onto the average firing rate axis in neural state space (**Fig. 4**). Similarly, a previous study ^25^ showed that across tasks, the population-response manifold rotates relative to the average firing rate axis. Although average firing rates differ, the underlying manifolds share a common geometry representing choice-related evidence. Thus, the averaged response dimension may obscure the latent structure of the population code, consistent with our present findings.

We have previously shown that both attention ^17^ and stimulus familiarity ^52^ enhance neural pattern separation. In this study, we extended these findings by demonstrating that attentional states in the fovea can enhance pattern separation in peripheral representations (**Fig. 5**). This result provides critical evidence for a functional coupling between foveal attentional engagement and peripheral visual processing during naturalistic search (see also ^16^). Importantly, the enhancement of pattern separation by foveal attention was not uniform but exhibited a non-linear profile, selectively amplifying target representations while compressing distractor representations (**Fig. 5**). This non-linear modulation suggests that attention does more than uniformly boost neural responses; rather, it restructures the representational geometry of peripheral visual populations to prioritize behaviorally relevant information. What neural mechanisms support the observed non-linear pattern separation? One possibility is that top-down inputs from prefrontal regions dynamically gate feature-specific pathways in visual cortex, selectively enhancing target-relevant signals ^53^. Another possibility is that local circuit mechanisms, such as inhibitory sharpening or normalization, interact with top-down signals to restructure population codes ^54^. Furthermore, our findings align with evidence that attention selectively reshapes the geometry of distributed semantic representations ^55^ and modulates neural representations to render reconstructions according to subjective appearance ^56^. Future studies are needed to further assess how attention changes the structure of the representational spaces over which it operates, including the spatial organization, feature maps, and object-based coding in the visual cortex ^57^ (see also ^17^).

In conclusion, our results reveal that goal-directed visual search emerges from dynamic, population-level coordination across cortical regions. By jointly encoding attention, category, memory, and spatial layout, neural populations flexibly transform and maintain task-relevant information throughout distinct stages of search. The discovery of orthogonal subspace representations further highlights how latent population geometry supports the encoding and maintenance of task-relevant information. These findings provide a unifying framework for understanding how distributed cortical networks orchestrate complex visual cognition. Future work should examine how these subspace dynamics are modulated by learning, context, and behavioral demands, and whether similar principles govern population coding in other goal-directed behaviors.

## Acknowledgements

This research was supported by Washington University in St. Louis. The funder had no role in study design, data collection and analysis, decision to publish, or preparation of the manuscript.

## Author Contributions

J.Z. designed research. J.Z. performed experiments. J.Z., N.R., and S.W. analyzed data. J.Z. and S.W. wrote the paper. All authors discussed the results and contributed toward the manuscript.

## Methods

### Data

We analyzed data from a publicly available dataset ^44^, in which recordings from areas V4, IT (including both TE and TEO), LPFC, and OFC were acquired from two male rhesus macaques weighing 12 and 15 kg.

### Tasks and stimuli

Monkeys were trained to perform a free-gaze visual search task. A central fixation point was presented for 400 ms, followed by a cue lasting 500 to 1300 ms. After a delay of 500 ms, the search array appeared. The search array contained 11 items, including two targets, which were randomly selected from a total of 20 predefined locations. Monkeys were required to find one of the two targets within 4000 ms and maintain fixation on the target for 800 ms to receive a juice reward. No constraints were placed on their search behavior, allowing the animals to perform the search naturally. Before the onset of the search array, monkeys were required to maintain central fixation. The two target stimuli belonged to the same category as the cue stimulus, though they were distinct images. There were four categories of stimuli—face, house, flower, and hand—each comprising 40 images. The cue stimulus was randomly selected from either the house or face category with equal probability. The remaining 9 stimuli in the search array were drawn from the other three categories. Each stimulus subtended an area of approximately 2° × 2°, with hue, saturation in the HSV color space, aspect ratio, and luminance matched across categories ^14,16,17^. The 20 locations, covering the visual field with eccentricities from 5° to 11°, included 18 locations symmetrically distributed across the left and right visual fields (9 on each side), and 2 locations on the vertical midline.

A visually guided saccade task was used to map the peripheral receptive fields (RFs) of recorded units. After a 400-ms central fixation, a stimulus (a face or house, identical to those in the visual search task) randomly appeared at one of 20 locations. Monkeys were required to make a saccade to the stimulus within 500 ms and maintain fixation on it for 300 ms to receive a reward.

Behavioral experiments were conducted using MonkeyLogic software (University of Chicago, IL), which presented stimuli, monitored eye movements, and triggered reward delivery.

### Electrophysiology

Single-unit and multi-unit spikes were recorded from V4, IT, LPFC, and OFC using 24– or 32-contact electrodes (V-Probe or S-Probe, Plexon Inc., Dallas, USA) with a 128-channel Cerebus System (Blackrock Microsystems, Salt Lake City, UT, USA). In most sessions, activity was recorded simultaneously from two of these areas. Neural recordings were filtered between 250 Hz and 5 kHz and digitized at 30 kHz to obtain spike data. Spike sorting was performed using Plexon’s Offline Sorter™ (OFS). LFP signals were obtained by filtering neural recordings between 0.3 and 250 Hz and digitizing them at 1000 Hz. Recording locations in V4, IT, LPFC, and OFC were verified using MRI. Eye movements were recorded with an infrared eye-tracking system (iViewX Hi-Speed, SensoMotoric Instruments [SMI], Teltow, Germany) at a sampling rate of 500 Hz.

### Spike rate

Measurements of neural activity were obtained from spike density functions, which were generated by convolving the time of action potentials with a function that projects activity forward in time (Growth = 1 ms, Decay = 20 ms) and approximates an EPSP ^58^. Specifically, this spike density function has two advantages ^58^. First, each spike exerts influence only forward in time, representing the actual postsynaptic effect of each cell. Second, by using a function that resembles a postsynaptic potential, we can apply time constants similar to those measured physiologically. Additionally, using the postsynaptic potential filter for time course analysis is advantageous because, when using the Gaussian filter, target discrimination times sometimes occur earlier than the unit’s evident visual latency. This impossible outcome occurs because, with the Gaussian filter, spikes exert influence backward in time. The spike rate of each unit was normalized by the mean baseline firing rate during the fixation spot preceding the cue.

### Receptive field

The visual response to the cue and the search array in the free-gaze visual search task was assessed by comparing the firing rate during the post-stimulus period (50 to 200 ms after cue/array onset) to the corresponding baseline (−150 to 0 ms relative to cue/array onset) using a Wilcoxon rank-sum test. Based on these responses, we classified units into three categories of RFs:

i. Units with a focal foveal RF: These units responded solely to the cue in the foveal region (P < 0.05) but not to the search array that included items in the periphery (P > 0.05).
ii. Units with a broad foveal RF: These units responded to both the cue and the search array (both Ps < 0.05).
iii. Units with a peripheral RF: These units only responded to the search array (P < 0.05) but not to the cue (P > 0.05). The RFs of these units were additionally mapped based on their activities in the visually guided saccade task. Units whose RFs could be mapped in this task had a localized peripheral RF, whereas units whose RFs could not be mapped had an unlocalized peripheral RF (i.e., units that responded to the search array onset but not during the saccade task; 11.87% of visually responsive units in V4, 5.58% in IT, 3.20% in OFC, and 58.29% in LPFC had an unlocalized RF).

Units not classified into the above categories (both Ps > 0.05) were not visually responsive and were excluded from further analysis.

### Selection of category-selective units

We selected *category-selective units* by comparing the response to face cues versus house cues in a time window of 50 to 200 ms after cue onset (Wilcoxon rank-sum test, P < 0.05). We further imposed a second criterion using a selectivity index similar to indices employed in previous IT studies ^59,60^. For each unit with a foveal RF, the response to face stimuli (*R*_face_) or house stimuli (*R*_house_) was calculated using the visual search task by subtracting the mean baseline activity (−150 to 0 ms relative to the onset of the cue) from the mean response to the face or house cue (50 to 200 ms after the onset of the cue). For each unit with a peripheral RF, *R*_face_ and *R*_house_ were calculated using the visually guided saccade task by subtracting the mean baseline activity (−150 to 0 ms relative to the peripheral stimulus onset) from the mean response to the saccade target (50 to 200 ms after the onset of the saccade target). It is worth noting that for both foveal and peripheral units, we ensured that there was only one stimulus in the RF. The selectivity index (SI) was then defined as (*R*_face_ − *R*_house_) / (*R*_face_ + *R*_house_). SI was set to 1 when *R*_face_ > 0 and *R*_house_ < 0, and to −1 when *R*_face_ < 0 and *R*_house_ > 0. Face-selective units were required to have an *R*_face_ at least 130% of *R*_house_ (i.e., the corresponding SI was greater than 0.13). Similarly, house-selective units were required to have an *R*_house_ at least 130% of *R*_face_ (i.e., the corresponding SI was smaller than −0.13). Units were labeled as non-category-selective if the response to face cues versus house cues was not significantly different (P > 0.05). The remaining units that did not fit into any of the aforementioned types were classified as undefined units (i.e., there was a significant difference but did not meet the second criterion). It is worth noting that we did not use the activity during the search to calculate the SI to minimize interactions with the attentional effect and between RFs.

### Selection of attention-selective units

We used the mean firing rate in a time window of 150 ms to 225 ms after fixation onset as the response to each fixation. For each unit, if there was a significant difference in response (determined using a two-tailed Wilcoxon signed-rank test, with a significance threshold of P < 0.05) between fixations on targets and fixations on distractors, it was classified as an *attention-selective unit*. Similarly, for units with a peripheral RF (as described above), we compared the response between targets and distractors within the RF in the same time window as for foveal units. Lastly, we calculated the attentional effect as the difference in firing rate between the same stimuli when they served as targets versus distractors.

### Angle between the category and attention hyperplanes

We used the method from ^32^ to measure the angle between the category and attention hyperplanes and examine their relationship. Specifically, we trained two classifiers: one to distinguish between faces and houses (category classifier; for distractor stimuli) and another to differentiate between target and distractor stimuli (attention classifier; for face and house stimuli) within the RF of units during the search. Classifiers were trained using the vector of firing rates (FRs) averaged over 0–255 ms from fixation onset for all foveal units within a region, following the same procedure, with the only difference being the condition groupings. This resulted in a matrix of mean FRs for each unit and each fixation (matrix size = units × fixations). We downsampled the data 100 times and trained a classifier on each sample. The final classifier was obtained by averaging the intercepts (*b*) and weights (**w**) of these 100 trained classifiers. The resulting classifier is a hyperplane defined by its orthogonal vector (size = the number of units) and an intercept. We calculated the angle between the two trained classifiers/hyperplanes as follows 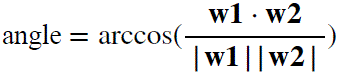: where **w1** and **w2** are the weight vectors of the two classifiers. To obtain the null distribution (1000 runs), we shuffled the condition labels for category and attention, trained two classifiers on the shuffled data, and calculated the angle between them to derive the shuffled angle.

### Principal component analysis (PCA)

When the activity of a population of units is plotted in a coordinate system in which each axis represents the firing rate of one unit—also known as the state space—the response dynamics of the population can be depicted as a high-dimensional neural trajectory. We used demixed PCA (dPCA; see below; **Fig. 2**) to compute a low-dimensional space and projected neural population activity onto the first three principal components (PCs) to visualize neural trajectories within the space of neural activity during the following periods (in 10 ms bins): (1) 0–500 ms from cue onset; (2) 0–500 ms from delay onset; and (3) 0–225 ms from fixation onset during search. The fixations/trials were sorted based on stimulus categories and/or attentional states. We further divided the neural population based on brain areas (V4, IT, OFC, and LPFC) and RFs (foveal and peripheral units). To compute statistics (i.e., identify time points along the trajectory that significantly differed between conditions), we shuffled the data between conditions 1000 times and compared the observed trajectory distances with those from the shuffled data. Multiple comparisons across time points were corrected using the false discovery rate (FDR; P < 0.05) ^61^.

PCA is an unsupervised method, and the resulting components can retain mixed information about category and attention coding. To better capture neural tuning to these two task parameters, we used dPCA, a dimensionality reduction technique that demixes the dependencies of population activity on task parameters by decomposing the data into a few components ^62^. Unlike PCA, the compression and decompression steps of dPCA do not directly reconstruct neural activity but instead reconstruct neural activity averaged over trials and some task parameters. We used dPCA with stimulus categories and/or attentional states and time as the marginalized variable. For the search period, the demixing weights (basis functions) were computed from the target responses. For the cue and delay periods, the demixing weights were computed from the averaged responses to faces and houses. Neural activity during each epoch was then projected onto the corresponding basis functions.

### Parallel axis and orthogonal plane

For **Fig. 4**, we first extracted trial-averaged PSTHs for each unit in response to eight face and eight house stimuli, from −60 to 210 ms relative to fixation onset, using 30-ms time bins. This resulted in nine time steps centered at [−45, −15, 15, 45, 75, 105, 135, 165, 195] ms. We selected 8 face and 8 house stimuli because, as the number of stimuli increases, the number of units that had seen all of them decreases. With 8 face and 8 house stimuli, approximately 41% to 73% of the units could be included in the analysis. Trials were grouped by stimulus identity and attentional status (target/distractor). We then concatenated each unit’s PSTHs into a response vector and combined the vectors of all units into a population response matrix with *T* × *C* rows and *N* columns, where *T* = 9 is the number of time steps, *C* = 32 is the number of conditions (16 stimulus identities × 2 attentional statuses), and *N* is the number of units. Only correct trials were used for the analyses. The neural population was divided based on brain region, RF, and selectivity. dPCA was performed across PSTHs for each subset of the population to decompose responses into two components, depending on either the attentional status task parameter (target and distractor components) or the category task parameter (face and house components), as described above. The neural population response for each condition at each time step can be depicted as a point in three-dimensional (3D) PC space. In this space, the projected population-averaged firing rate changed along a linear axis. A vector corresponding to this linear projection axis can be readily derived from the PC coefficients ^25^: 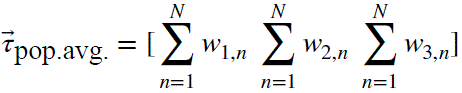, where 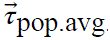. is the axis of the population average firing rate, and *w_i,n_* is the coefficient of unit *n* for the *i*-th PC.

We projected each point onto the axis of the average firing rate in the 3D PC space. The projection of a point *P* onto the average FR axis is: 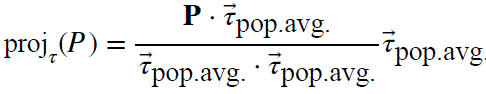. The parallel components along the average FR axis is determined by the Euclidean distance between the projected point proj*_τ_*(*P*) and the origin, with the sign corresponding to the sign of the dot product 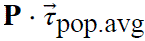.

The normal vector to a plane orthogonal to the population average FR axis is 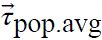. We define the orthogonal plane as the plane that passes through the origin. The unit vector along the *y*-axis of the plane is determined as: 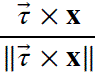, where **x** is the vector along the *x*-axis in the plane, which can be obtained by projecting an arbitrary point in the 3D space, with the specific point [10, 0, 0] used in this case. The projection of a point *P* onto the plane is: 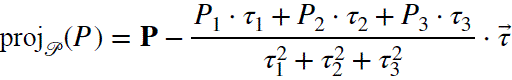. We then projected 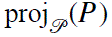 again onto the *x*-axis and *y*-axis of the plane to obtain the coordinates in the plane. These coordinates are determined by the Euclidean distances between the projected points and the origin, with the sign corresponding to the sign of the dot product, respectively. The orthogonal components to the average FR axis are defined by these coordinates.

We used linear discriminant analysis (LDA) to classify the target stimulus from the distractor or the face stimulus from the house. The classifier used parallel and orthogonal components for classification, respectively.

### Geometry in neural subspaces using rotated PCA (rPCA)

For each subspace of the neural state space, standard PCA was performed across different conditions to identify the first two or three axes that captured the most response variance due to condition variation in each subspace. To compare representational geometries of different subspaces, the subspace axes were rotated and scaled (i.e., rPCA procedure) to align them (see details in ^37^).

We sorted/reconstructed the neural responses as follows:

For **Fig. 4**, neural responses of foveal units from V4 and IT were sorted according to the stage of visual search and stimulus identity, and only stimuli presented to all neurons were included. The numbers of shared face and house stimuli for the V4 and IT populations were 32/36 and 27/30, respectively. The time windows corresponding to the different task stages were defined as follows: 0–500 ms from cue onset, 0–500 ms from delay onset, and 0–250 ms from fixation onset during the search period.

For **Fig. 5**, we used a multivariable linear regression model to assess how the categories and attentional states of multiple stimuli within the RF of a peripheral unit, along with foveal content and attentional state during search, collectively shaped the neural response, because the RF of a peripheral unit may contain multiple stimuli. This model allowed us to dissociate the contributions of each variable for subsequent modeling. Task variables were defined by stimulus category and attentional state. Specifically, RF stimuli included face target, face distractor, house target, house distractor, flower distractor, and hand distractor, crossed with fixation type (fixation on a target versus fixation on a distractor), resulting in a 12-dimensional one-hot vector representation. For example, the vector (1 0 0 0 1 1 0 0 0 0 0 0)^T^ denotes a fixation on a target with a face target, flower distractor, and hand distractor in the peripheral RF. In the model, the average neural response during each fixation (0–250 ms after fixation onset) was modeled as a linear combination of these task variables. To prevent overfitting, we applied Lasso regularization, with the regularization strength determined by maximum-likelihood estimation. For each unit, fixations were randomly split in half 100 times, and the regression model was fitted separately to each half. This procedure produced 200 estimates of each regression coefficient, which were then averaged. Subsequent analyses were performed on these mean coefficient values.

For **Fig. 6**, neural responses were sorted into four conditions (fixations on a face, house, flower, or hand stimulus) × two groups (fixating on the stimulus for the first time in each trial or re-fixating). There were three subsets of populations (foveal units from V4, IT, and OFC).

For **Fig. 7**, neural responses of foveal units in V4 and IT from 0 to 200 ms after fixation onset were grouped into 20 conditions corresponding to the 20 spatial fixation locations. To compare representational geometries across time, the responses were further divided into 50-ms time bins. Neural activity within each time bin defined a subspace. These four subspaces were rotated and scaled as described above and further aligned according to the *x*–*y* spatial coordinates of the fixation locations.

We used time-averaged responses (for **Fig. 3** and **Fig. 6**) or regression coefficients (for **Fig. 5**) to construct neuronal vectors and calculated the Euclidean distance between them to quantify neural representational distance. Changes in population geometry were further characterized using the cosine angle between neuronal vectors: 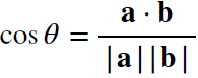, where **a** and **b** represent neuronal vectors for different conditions.

### Representational similarity analysis (RSA)

For RSA ^63^, dissimilarity matrices (DMs) are symmetrical matrices representing dissimilarity between all pairs of locations. In a DM, larger values indicate greater dissimilarity (distance) between pairs, with the smallest possible value (0) indicating similarity of a condition to itself. We used Pearson correlation to compute DMs for neural populations and Euclidean distance for stimulus location coordinates on the screen. Spearman’s correlation was used to assess the correspondence between DMs, as it does not assume a linear relationship ^64^. Specifically, PSTHs of the foveal units from 0 to 200 ms after fixation onset (with a 50 ms time bin) were sorted based on the location of the fixated stimulus. The average response for each of the four time steps was used to compute the dissimilarity value (1 − Pearson’s *r*) for the neural DM between each pair of locations. To assess the significance of the correspondence between the neural and physical DMs, we employed permutation tests with 1000 runs. In each run, location labels were randomly shuffled, and the correlation between DMs was recalculated. The distribution of correlation coefficients from the shuffled data (i.e., the null distribution; shaded in gray in **Fig. 7**) was then compared to the observed correlation coefficient (i.e., unshuffled; connected dots in **Fig. 7**). If the observed correlation exceeded 95% of the coefficients in the null distribution, it was deemed significant. Bonferroni correction was applied to account for multiple comparisons.

A similar procedure was applied to examine the relationships between different stages of the visual search process (**Fig. 4**).

### Code availability

The source code for this study is publicly available on OSF (https://osf.io/sdgkr/).

**Fig. S1.**
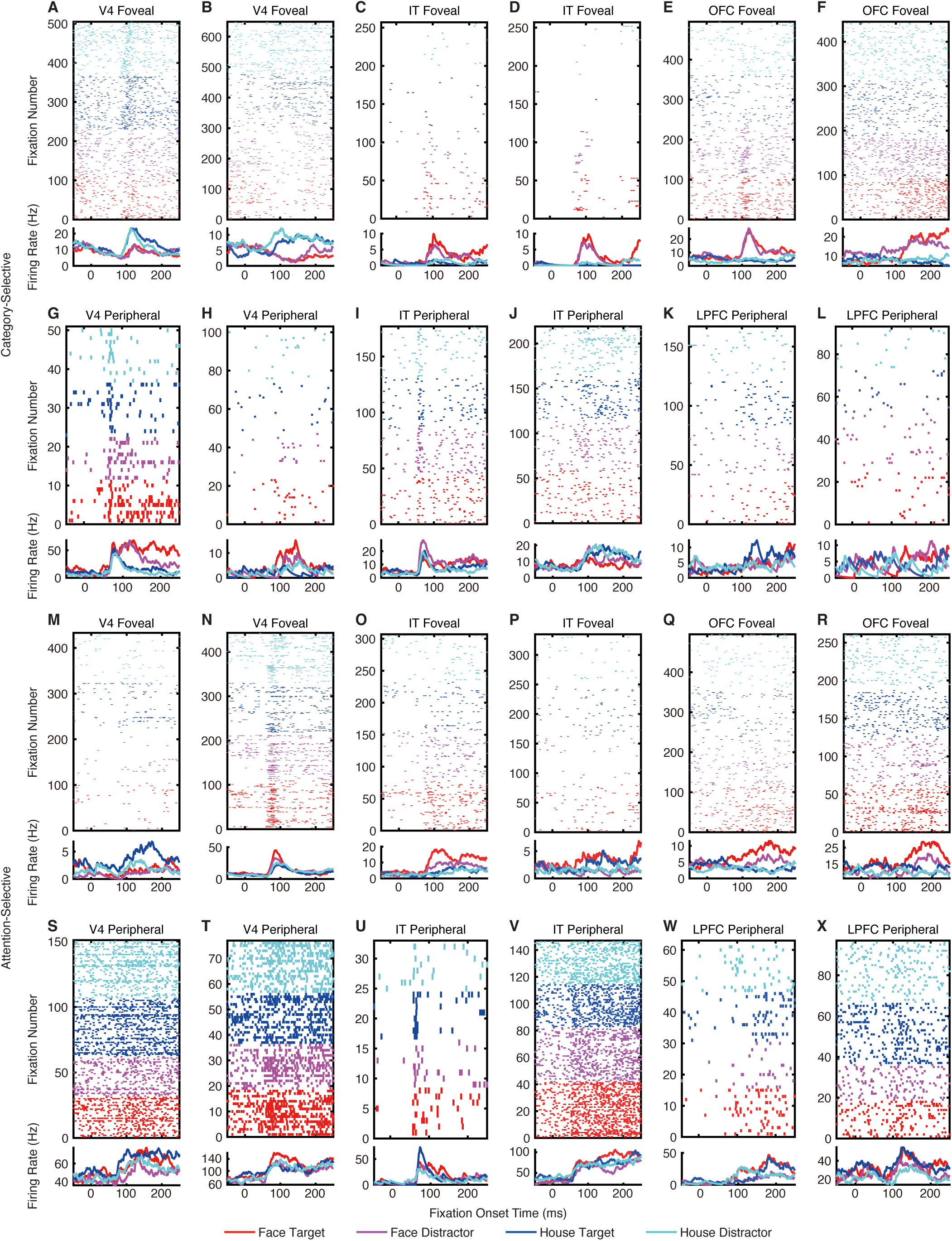
Additional results for category-selective and attention-selective units. (**A–F**) Category-selective foveal units. **(G–L)** Category-selective peripheral units. **(M–R)** Attention-selective foveal units. **(S–X)** Attention-selective peripheral units. **(A, B, G, H, M, N, S, T)** V4 units. **(C, D, I, J, O, P, U, V)** IT units. **(E, F, Q, R)** OFC units. **(K, L, W, X)** LPFC units. Legend conventions as in Fig. 1.

**Fig. S2.**
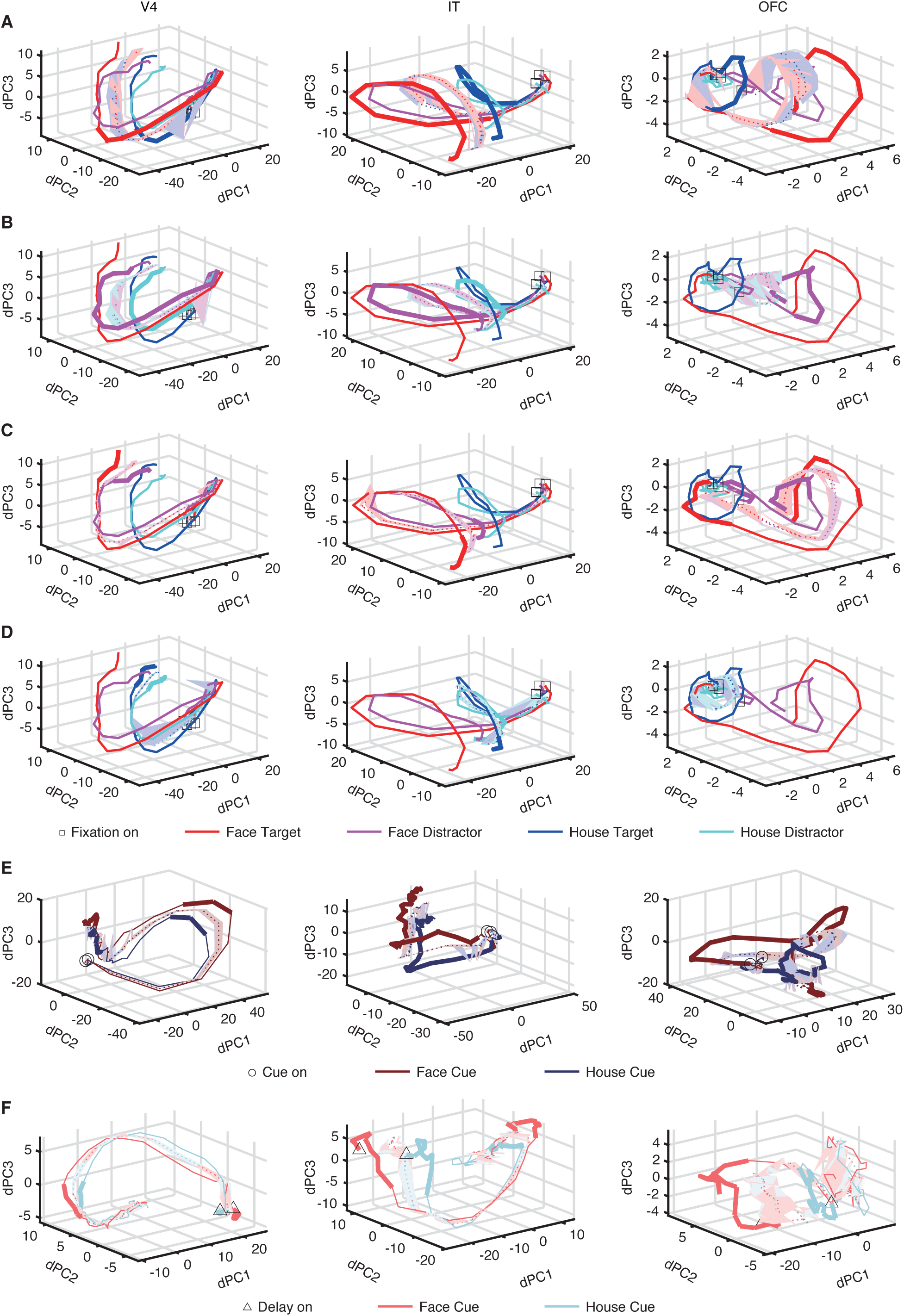
Control results for neural population dynamics. (**A–D**) Statistical comparison using permutation tests for each brain area in the main (category-matching) task. **(A)** Comparison of face targets versus house targets. **(B)** Comparison of face versus house distractors. **(C)** Comparison of face targets versus face distractors. **(D)** Comparison of house targets versus house distractors. **(E, F)** Replication of results in the identity-matching task. In this task, there was only one search target, and monkeys were required to fixate on the identical target matching the cue. Legend conventions as in Fig. 2.

**Fig. S3.**
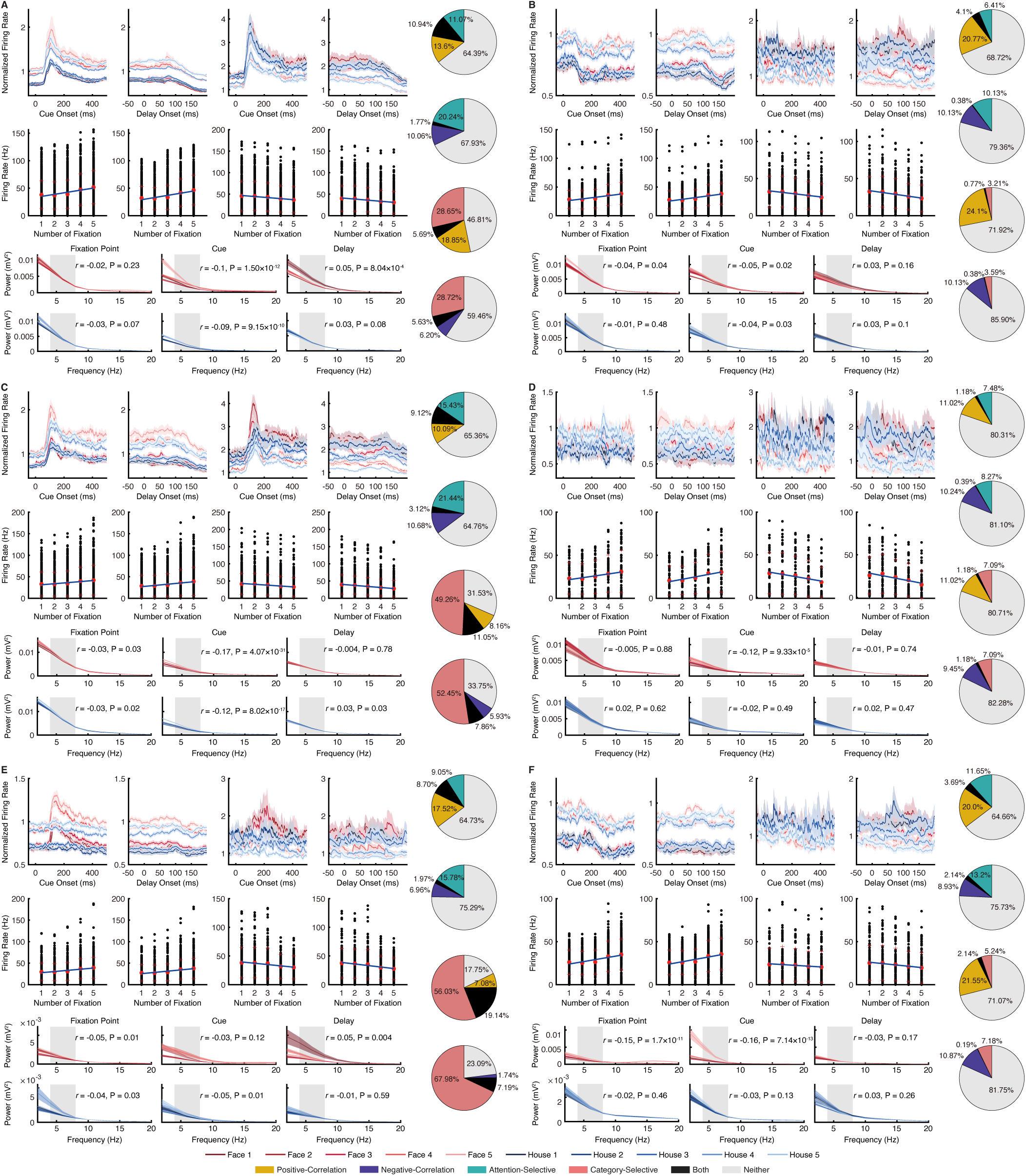
Neural activity during cue presentation and maintenance, analyzed separately for foveal and peripheral units. **(A)** V4 foveal units. **(B)** V4 peripheral units. **(C)** IT foveal units. **(D)** IT peripheral units. **(E)** OFC foveal units. **(F)** LPFC peripheral units. Legend conventions as in Fig. 3.

**Fig. S4.**
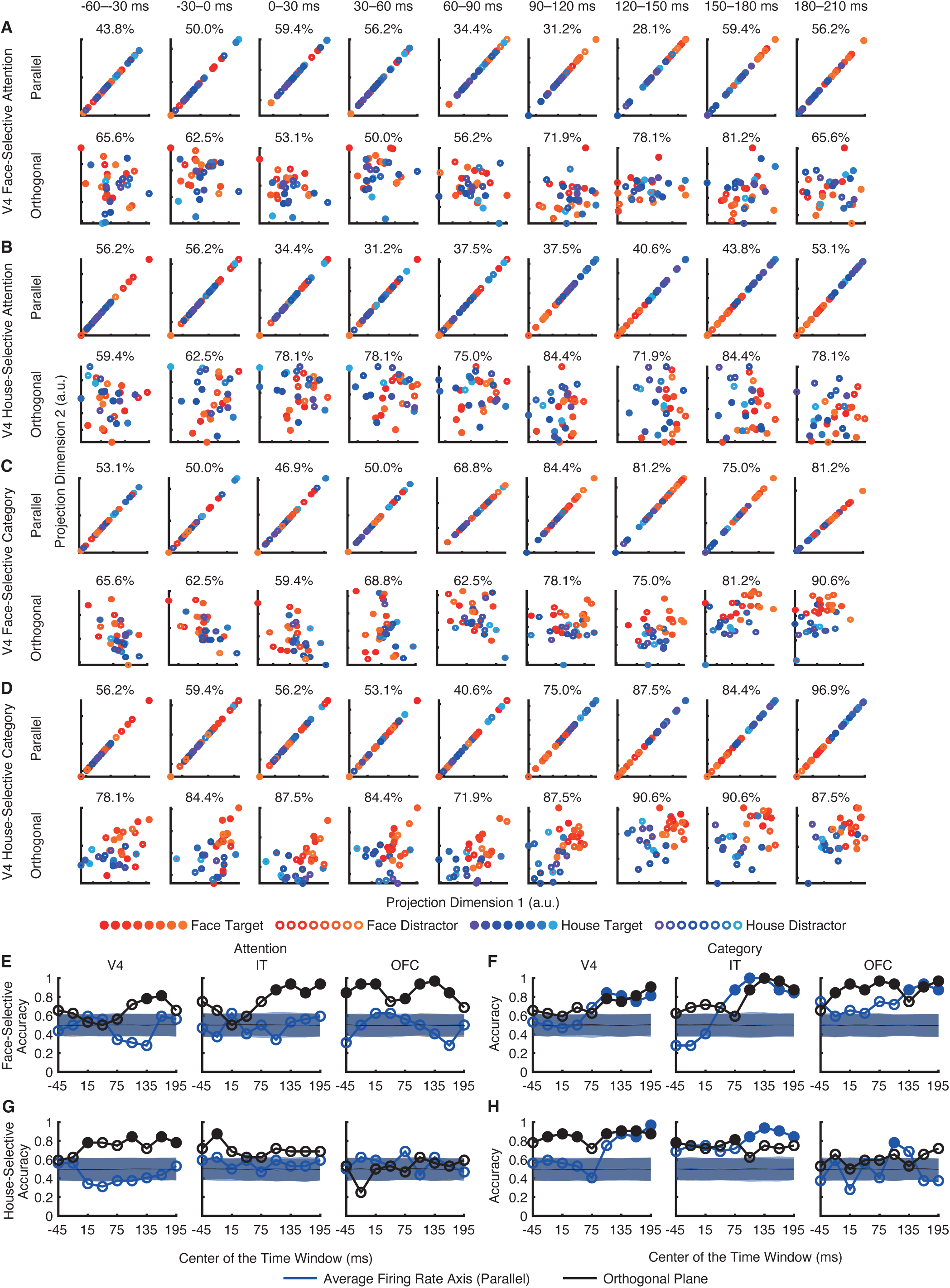
Neural processing across stages of visual search in the orthogonal subspace for face-selective and house-selective units. Legend conventions as in Fig. 4.

**Fig. S5.**
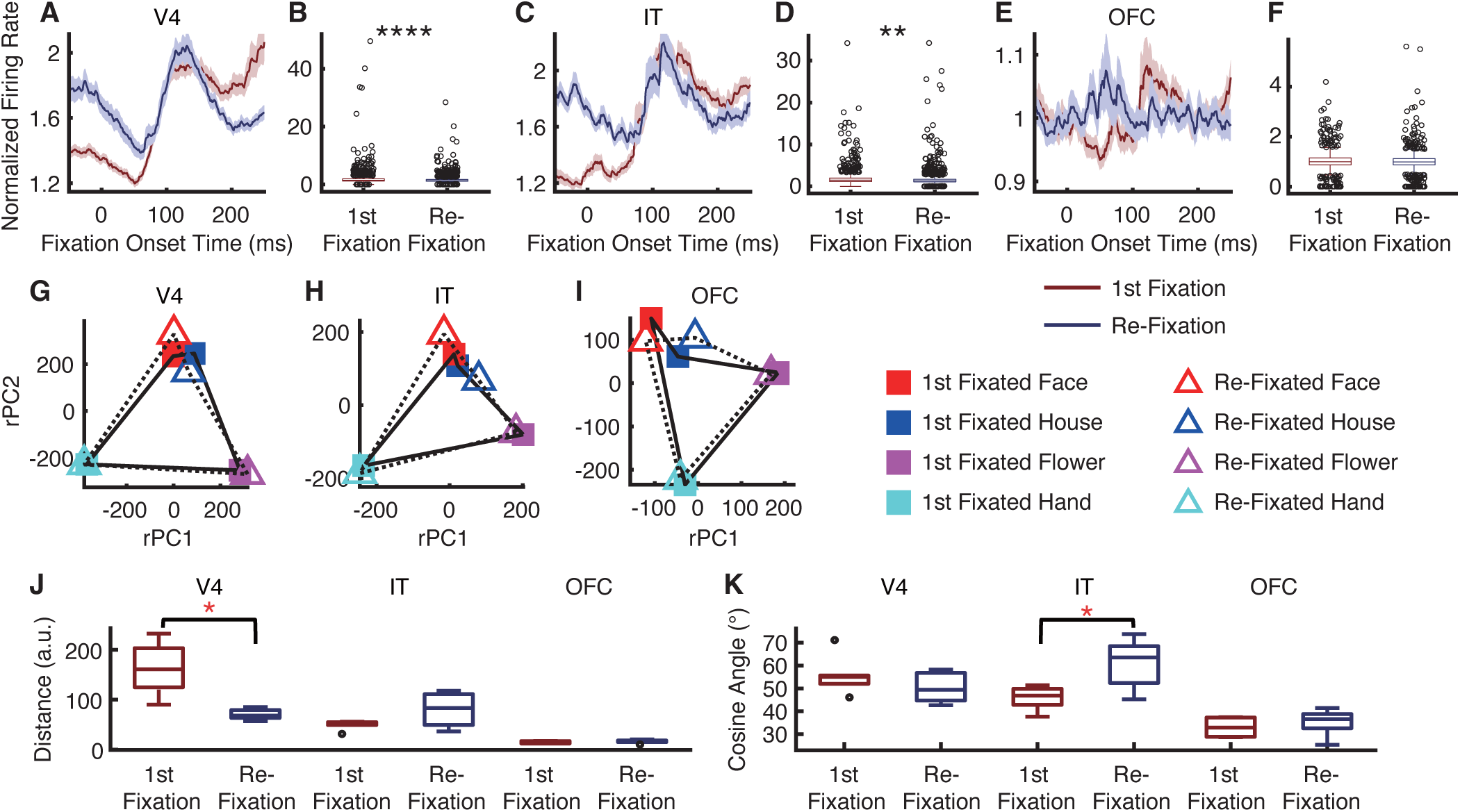
Comparison of first-fixated versus re-fixated search items using only distractor stimuli. Legend conventions as in Fig. 6.

## References

1. Kastner, S., and Ungerleider, L.G. (2000). Mechanisms of visual attention in the human cortex. Annual Review of Neuroscience.

2. Corbetta, M., and Shulman, G.L. (2002). Control of goal-directed and stimulus-driven attention in the brain. Nat Rev Neurosci 3, 201–215.

3. Petersen, S.E., and Posner, M.I. (2012). The Attention System of the Human Brain: 20 Years After. Annual Review of Neuroscience 35, 73–89. 10.1146/annurev-neuro-062111-150525.

4. Chelazzi, L., Miller, E.K., Duncan, J., and Desimone, R. (1993). A neural basis for visual search in inferior temporal cortex. Nature 363, 345–347.

5. Buschman, T.J., and Miller, E.K. (2007). Top-Down Versus Bottom-Up Control of Attention in the Prefrontal and Posterior Parietal Cortices. Science 315, 1860–1862. doi:10.1126/science.1138071.

6. Buschman, Timothy J., and Kastner, S. (2015). From Behavior to Neural Dynamics: An Integrated Theory of Attention. Neuron 88, 127–144. 10.1016/j.neuron.2015.09.017.

7. Zhou, H., Schafer, Robert J., and Desimone, R. (2016). Pulvinar-Cortex Interactions in Vision and Attention. Neuron 89, 209–220. 10.1016/j.neuron.2015.11.034.

8. Moore, T., and Zirnsak, M. (2017). Neural Mechanisms of Selective Visual Attention. Annual Review of Psychology 68, 47–72. 10.1146/annurev-psych-122414-033400.

9. Fiebelkorn, I.C., and Kastner, S. (2020). Functional Specialization in the Attention Network. Annual Review of Psychology 71, 221–249. 10.1146/annurev-psych-010418-103429.

10. Sheinberg, D.L., and Logothetis, N.K. (2001). Noticing Familiar Objects in Real World Scenes: The Role of Temporal Cortical Neurons in Natural Vision. The Journal of Neuroscience 21, 1340–1350.

11. Monosov, I.E., Sheinberg, D.L., and Thompson, K.G. (2010). Paired neuron recordings in the prefrontal and inferotemporal cortices reveal that spatial selection precedes object identification during visual search. Proceedings of the National Academy of Sciences 107, 13105–13110. 10.1073/pnas.1002870107.

12. Bichot, Narcisse P., Heard, Matthew T., DeGennaro, Ellen M., and Desimone, R. (2015). A Source for Feature-Based Attention in the Prefrontal Cortex. Neuron 88, 832–844. 10.1016/j.neuron.2015.10.001.

13. Stemmann, H., and Freiwald, W.A. (2019). Evidence for an attentional priority map in inferotemporal cortex. Proceedings of the National Academy of Sciences 116, 23797–23805. doi:10.1073/pnas.1821866116.

14. Zhang, J., Zhu, X., Zhou, H., and Wang, S. (2024). Behavioral and Neural Mechanisms of Face-Specific Attention during Goal-Directed Visual Search. The Journal of Neuroscience 44, e1299242024. 10.1523/JNEUROSCI.1299-24.2024.

15. Saalmann, Y.B., Pinsk, M.A., Wang, L., Li, X., and Kastner, S. (2012). The Pulvinar Regulates Information Transmission Between Cortical Areas Based on Attention Demands. Science 337, 753–756. doi:10.1126/science.1223082.

16. Zhang, J., Zhou, H., and Wang, S. (2024). Distinct visual processing networks for foveal and peripheral visual fields. Communications Biology 7, 1259. 10.1038/s42003-024-06980-2.

17. Zhang, J., Cao, R., Zhu, X., Zhou, H., and Wang, S. (2025). Distinct attentional characteristics of neurons with visual feature coding in the primate brain. Science Advances 11, eadq0332. doi:10.1126/sciadv.adq0332.

18. Zhang, Y., Meyers, E.M., Bichot, N.P., Serre, T., Poggio, T.A., and Desimone, R. (2011). Object decoding with attention in inferior temporal cortex. Proceedings of the National Academy of Sciences 108, 8850–8855. 10.1073/pnas.1100999108.

19. Logothetis, N.K., and Sheinberg, D.L. (1996). Visual object recognition. Annual Review of Neuroscience 19, 577–621. doi:10.1146/annurev.ne.19.030196.003045.

20. Gross, C.G. (1994). How Inferior Temporal Cortex Became a Visual Area. Cerebral Cortex 4, 455–469. 10.1093/cercor/4.5.455.

21. Tanaka, K. (1997). Mechanisms of visual object recognition: monkey and human studies. Current Opinion in Neurobiology 7, 523–529. 10.1016/S0959-4388(97)80032-3.

22. Tsao, D.Y., Freiwald, W.A., Tootell, R.B.H., and Livingstone, M.S. (2006). A Cortical Region Consisting Entirely of Face-Selective Cells. Science 311, 670–674. 10.1126/science.1119983.

23. Bao, P., She, L., McGill, M., and Tsao, D.Y. (2020). A map of object space in primate inferotemporal cortex. Nature 583, 103–108. 10.1038/s41586-020-2350-5.

24. Wolfe, J.M. (2021). Guided Search 6.0: An updated model of visual search. Psychonomic Bulletin & Review 28, 1060–1092. 10.3758/s13423-020-01859-9.

25. Okazawa, G., Hatch, C.E., Mancoo, A., Machens, C.K., and Kiani, R. (2021). Representational geometry of perceptual decisions in the monkey parietal cortex. Cell 184, 3748–3761.e3718. 10.1016/j.cell.2021.05.022.

26. Churchland, M.M., Cunningham, J.P., Kaufman, M.T., Foster, J.D., Nuyujukian, P., Ryu, S.I., and Shenoy, K.V. (2012). Neural population dynamics during reaching. Nature 487, 51–56. 10.1038/nature11129.

27. Sun, X., O’Shea, D.J., Golub, M.D., Trautmann, E.M., Vyas, S., Ryu, S.I., and Shenoy, K.V. (2022). Cortical preparatory activity indexes learned motor memories. Nature 602, 274–279. 10.1038/s41586-021-04329-x.

28. Safaie, M., Chang, J.C., Park, J., Miller, L.E., Dudman, J.T., Perich, M.G., and Gallego, J.A. (2023). Preserved neural dynamics across animals performing similar behaviour. Nature 623, 765–771. 10.1038/s41586-023-06714-0.

29. Chaisangmongkon, W., Swaminathan, S.K., Freedman, D.J., and Wang, X.J. (2017). Computing by Robust Transience: How the Fronto-Parietal Network Performs Sequential, Category-Based Decisions. Neuron 93, 1504–1517 e1504. 10.1016/j.neuron.2017.03.002.

30. Aoi, M.C., Mante, V., and Pillow, J.W. (2020). Prefrontal cortex exhibits multidimensional dynamic encoding during decision-making. Nat Neurosci 23, 1410–1420. 10.1038/s41593-020-0696-5.

31. Minxha, J., Adolphs, R., Fusi, S., Mamelak, A.N., and Rutishauser, U. (2020). Flexible recruitment of memory-based choice representations by the human medial frontal cortex. Science 368. 10.1126/science.aba3313.

32. Libby, A., and Buschman, T.J. (2021). Rotational dynamics reduce interference between sensory and memory representations. Nat Neurosci 24, 715–726. 10.1038/s41593-021-00821-9.

33. Nieh, E.H., Schottdorf, M., Freeman, N.W., Low, R.J., Lewallen, S., Koay, S.A., Pinto, L., Gauthier, J.L., Brody, C.D., and Tank, D.W. (2021). Geometry of abstract learned knowledge in the hippocampus. Nature 595, 80–84. 10.1038/s41586-021-03652-7.

34. Fu, Z., Beam, D., Chung, J.M., Reed, C.M., Mamelak, A.N., Adolphs, R., and Rutishauser, U. (2022). The geometry of domain-general performance monitoring in the human medial frontal cortex. Science 376, eabm9922. 10.1126/science.abm9922.

35. Liu, C., Jia, S., Liu, H., Zhao, X., Li, C.T., Xu, B., and Zhang, T. (2025). Recurrent neural networks with transient trajectory explain working memory encoding mechanisms. Commun Biol 8, 137. 10.1038/s42003-024-07282-3.

36. Sohn, H., Narain, D., Meirhaeghe, N., and Jazayeri, M. (2019). Bayesian Computation through Cortical Latent Dynamics. Neuron 103, 934–947.e935. 10.1016/j.neuron.2019.06.012.

37. Xie, Y., Hu, P., Li, J., Chen, J., Song, W., Wang, X.-J., Yang, T., Dehaene, S., Tang, S., Min, B., and Wang, L. (2022). Geometry of sequence working memory in macaque prefrontal cortex. Science 375, 632–639. 10.1126/science.abm0204.

38. Mohan, K., Zhu, O., and Freedman, D.J. (2021). Interaction between neuronal encoding and population dynamics during categorization task switching in parietal cortex. Neuron 109, 700–712 e704. 10.1016/j.neuron.2020.11.022.

39. Ni, A.M., Ruff, D.A., Alberts, J.J., Symmonds, J., and Cohen, M.R. (2018). Learning and attention reveal a general relationship between population activity and behavior. Science 359, 463–465. 10.1126/science.aao0284.

40. Mante, V., Sussillo, D., Shenoy, K.V., and Newsome, W.T. (2013). Context-dependent computation by recurrent dynamics in prefrontal cortex. Nature 503, 78–84. 10.1038/nature12742.

41. Langdon, C., and Engel, T.A. (2025). Latent circuit inference from heterogeneous neural responses during cognitive tasks. Nat Neurosci 28, 665–675. 10.1038/s41593-025-01869-7.

42. Bichot, N.P., Rossi, A.F., and Desimone, R. (2005). Parallel and Serial Neural Mechanisms for Visual Search in Macaque Area V4. Science 308, 529–534. 10.1126/science.1109676.

43. Zhou, H., and Desimone, R. (2011). Feature-Based Attention in the Frontal Eye Field and Area V4 during Visual Search. Neuron 70, 1205–1217. 10.1016/j.neuron.2011.04.032.

44. Zhang, J., Zhu, X., Zhou, H., and Wang, S. (2025). A large neuronal dataset for natural category-based free-gaze visual search in macaques. Scientific Data 12, 779. 10.1038/s41597-025-05130-5.

45. Wang, S., Mamelak, A.N., Adolphs, R., and Rutishauser, U. (2018). Encoding of Target Detection during Visual Search by Single Neurons in the Human Brain. Current Biology 28, 2058–2069.e2054. 10.1016/j.cub.2018.04.092.

46. Gothard, K.M. (2020). Multidimensional processing in the amygdala. Nature Reviews Neuroscience 21, 565–575. 10.1038/s41583-020-0350-y.

47. Mazer, J.A., and Gallant, J.L. (2003). Goal-Related Activity in V4 during Free Viewing Visual Search: Evidence for a Ventral Stream Visual Salience Map. Neuron 40, 1241–1250. 10.1016/S0896-6273(03)00764-5.

48. Zhang, J., Xia, J., Zhou, H., and Wang, S. (2025). Gamma synchronization between the medial temporal lobe and medial frontal cortex for goal-directed visual attention in humans. Cell Reports 44. 10.1016/j.celrep.2025.115905.

49. D’Esposito, M. (2007). From cognitive to neural models of working memory. Philosophical Transactions of the Royal Society B: Biological Sciences 362, 761–772. doi:10.1098/rstb.2007.2086.

50. Kaminski, J., Sullivan, S., Chung, J.M., Ross, I.B., Mamelak, A.N., and Rutishauser, U. (2017). Persistently active neurons in human medial frontal and medial temporal lobe support working memory. Nat Neurosci 20, 590–601. 10.1038/nn.4509 http://www.nature.com/neuro/journal/v20/n4/abs/nn.4509.html#supplementary-information.

51. Siegel, M., Warden, M.R., and Miller, E.K. (2009). Phase-dependent neuronal coding of objects in short-term memory. Proceedings of the National Academy of Sciences 106, 21341–21346. doi:10.1073/pnas.0908193106.

52. Cao, R., Wang, J., Brunner, P., Willie, J.T., Li, X., Rutishauser, U., Brandmeir, N.J., and Wang, S. (2024). Neural mechanisms of face familiarity and learning in the human amygdala and hippocampus. Cell Reports 43, 113520. 10.1016/j.celrep.2023.113520.

53. Miller, E.K., and Cohen, J.D. (2001). An Integrative Theory of Prefrontal Cortex Function. Annual Review of Neuroscience 24, 167–202. 10.1146/annurev.neuro.24.1.167.

54. Reynolds, J.H., and Heeger, D.J. (2009). The Normalization Model of Attention. Neuron 61, 168–185. 10.1016/j.neuron.2009.01.002.

55. Nastase, S.A., Connolly, A.C., Oosterhof, N.N., Halchenko, Y.O., Guntupalli, J.S., Visconti di Oleggio Castello, M., Gors, J., Gobbini, M.I., and Haxby, J.V. (2017). Attention Selectively Reshapes the Geometry of Distributed Semantic Representation. Cerebral Cortex 27, 4277–4291. 10.1093/cercor/bhx138.

56. Horikawa, T., and Kamitani, Y. (2022). Attention modulates neural representation to render reconstructions according to subjective appearance. Communications Biology 5, 34. 10.1038/s42003-021-02975-5.

57. Chapman, A.F., and Störmer, V.S. (2024). Representational structures as a unifying framework for attention. Trends in Cognitive Sciences 28, 416–427. 10.1016/j.tics.2024.01.002.

58. Thompson, K.G., Hanes, D.P., Bichot, N.P., and Schall, J.D. (1996). Perceptual and motor processing stages identified in the activity of macaque frontal eye field neurons during visual search. Journal of Neurophysiology 76, 4040–4055.

59. Freiwald, W.A., Tsao, D.Y., and Livingstone, M.S. (2009). A face feature space in the macaque temporal lobe. Nat Neurosci 12, 1187–1196. http://www.nature.com/neuro/journal/v12/n9/suppinfo/nn.2363_S1.html.

60. Freiwald, W.A., and Tsao, D.Y. (2010). Functional Compartmentalization and Viewpoint Generalization Within the Macaque Face-Processing System. Science 330, 845. 10.1126/science.1194908.

61. Benjamini, Y., and Hochberg, Y. (1995). Controlling the False Discovery Rate: A Practical and Powerful Approach to Multiple Testing. Journal of the Royal Statistical Society. Series B (Methodological) 57, 289–300.

62. Kobak, D., Brendel, W., Constantinidis, C., Feierstein, C.E., Kepecs, A., Mainen, Z.F., Qi, X.L., Romo, R., Uchida, N., and Machens, C.K. (2016). Demixed principal component analysis of neural population data. Elife 5. 10.7554/eLife.10989.

63. Kriegeskorte, N., Mur, M., and Bandettini, P. (2008). Representational similarity analysis - connecting the branches of systems neuroscience. Frontiers in Systems Neuroscience 2. 10.3389/neuro.06.004.2008.

64. Stolier, R.M., and Freeman, J.B. (2016). Neural pattern similarity reveals the inherent intersection of social categories. Nat Neurosci 19, 795–797. 10.1038/nn.4296 http://www.nature.com/neuro/journal/v19/n6/abs/nn.4296.html#supplementary-information.

